# Chemically induced *in vivo* retinal neuron regeneration rescues vision in mice

**DOI:** 10.1101/2023.12.27.572921

**Authors:** Rana Muhammad Shoaib, Sasha Medvidovik, Laura Korobkova, Pratigya Tripathi, Ryan P. Lin, Aregnazan Sandrosyan, Swaminathan Hariharan, Biraj Mahato

## Abstract

Loss of retinal ganglion cells (RGCs) is a major cause of vision loss in optic neuropathies such as glaucoma, with no available treatments to restore vision. In teleost fish, Müller glia possesses a remarkable regenerative capacity to replace lost RGCs and restore vision—a capability lacking in mammals. Here, we have identified a six-small molecule cocktail (6C) that induces *in vivo* reprogramming of retina resident Müller glia into retinal neurons within the ganglion cell layer (GCL) following RGC injury. We name these cells “chemically induced GCL neurons (CiGN)”. During reprogramming process, Müller glia re-enters the cell cycle in the inner nuclear layer, asymmetrically divide, proliferate and migrate to the GCL as SOX2^+^ and SOX2^−^ intermediates, exit the cell cycle, and differentiate into CiGN cells—mirroring some aspects of retinal regeneration seen in teleost fish. Functionally, 6C treatment restores long-term visual functions in rodent models of ocular hypertension and NMDA-induced RGC injury. Notably, 6C induces axon extension along the optic nerve and establish connections to the lateral geniculate nucleus (LGN) possibly through a neuronal relay mechanism. These findings highlight small molecule mediated cellular reprogramming as a potential therapeutic strategy for vision restoration in glaucoma and other optic neuropathies that affects millions of children and adults worldwide.

## INTRODUCTION

The regenerative capacity of the adult mammalian central nervous system (CNS), including the neurosensory retina, is limited ^1–3^. However, recent research has shown that Müller glia, a type of support cell in the retina, can be reprogrammed into retinal neurons *in vivo* ^4–9^. In lower vertebrates, Müller glia naturally transforms into neural progenitor cells and generate functional neurons to restore vision after injury. Researchers are currently addressing the limitation of endogenous mammalian retinal neurogenesis in two main ways: i) transplanting *in vitro* engineered stem cell derived retinal neurons or neural progenitor cells into injured retina^10–16^ or ii) overexpression of transcription factors in Müller cells to regenerate neural progenitors or retinal neurons to restore retinal function^6, 17–20^. However, these methods involve the use of viruses which may present challenges for its clinical implementation^9, 21^. An alternative method, using small molecules for chemical reprogramming, has recently garnered considerable attention ^22–25^. A prior study demonstrated that a combination of five small molecules (5C) could reprogram fibroblasts into photoreceptors like cells within 10 days, subsequently leading to vision restoration upon subretinal transplantation in blind mice^26^. Molecular analysis identified ASCL1 as crucial for photoreceptor like reprogramming, with its role in promoting retinal neurogenesis from Müller glia also supported by other studies ^17, 19, 20, 27–29^. Recent research shows small molecules can reprogram glial cells into functional neurons and fibroblasts into functional cardiac cells *in vivo*^30, 31^. Based on this, we hypothesized that modifying our cocktail could induce Müller cells to become retinal ganglion like neurons *in vivo*. Modifying this combination with additional molecules supplementary molecules, we report a novel chemical combination (6C) that stimulates reprogramming of resident retinal Müller cells into CiGNs and rescues vision in DBA/2J and NMDA-injured mice where RGC injury causes vision loss. To validate the Müller origin of CiGNs, we conducted lineage tracing experiments. Moreover, newly generated CiGN cells project axons to optic nerve (ON) and establish connections lateral geniculate nucleus (LGN) which supports visual function rescue.

## RESULTS

### 6C induces *in vivo* Müller glia reprogramming to CiGN in 12.5-mo-old DBA/2J mice

During 5C mediated photoreceptor-like cell reprogramming, we observed emergence of cells with morphological similarities to RGCs between days 4 and 5^26^. This prompted us to hypothesize that RGC generation from fibroblasts might occur earlier in this reprogramming process in the same dish. Using an Sncg promoter reporter^32^, we detected GFP^+^ cells as early as day 4 during fibroblast reprogramming, suggesting an RGC lineage commitment, albeit at a low frequency. We also identified PAX6^+^ cells (a progenitor marker) on day 3, suggesting that 5C mediated reprogramming of human fibroblasts into retinal neurons may proceed through a progenitor-like stage^33^. We next asked whether 5C could reprogram human Müller cells into chemically induced RGC like neurons (CiRGCs). Encouragingly, we observed Sncg-driven GFP expression in Müller-derived CiRGCs. To improve efficiency, we screened additional molecules and identified a new five chemical combination that reprograms human Muller cells into CiRGCs *in vitro* (**Figure S1**). scRNA-Seq analysis of these cells revealed clusters closely aligned with native RGCs from day 59 fetal retina in published studies (**Figure S2**)^34, 35^. These clusters expressed all canonical RGC markers, confirming their commitment to the RGC like neuronal lineage.

To investigate whether this five-chemical combination could reprogram endogenous Müller cells into CiGN cells, we intravitreally injected the cocktail into DBA/2J mice (12-month-old) and assessed Müller cell cycle entry using Bromodeoxyuridine (BrdU) uptake. We detected BrdU^+^ cells in the INL layer by day 1, albeit at low frequency. At this stage, we refined our cocktail by screening additional small molecules and identified a six-chemical combination [valproic acid (V), CHIR99021(C), RepSox (R), Forskolin(F) (base molecules); and IWR1, S24795] supplemented with two factors, Noggin and IGF1 (referred to as 6C plus 2F), which significantly enhanced Müller glia cell cycle entry by day 1. Next we intravitreally injected 6C into the eyes of 12-month-old DBA/2J mice, a stage characterized by peak ocular hypertension and significant RGC loss, as validated by pattern ERG and tracked the fate of the BrdU^+^ Muller cells^36^. The retinas were isolated and examined on days 1, 2, 3 and 6 after 6C intravitreal injections and these timepoints are selected based on prior publications^37, 38^. We observed BrdU^+^ cells along the inner nuclear layer (INL) on day 1, subsequent migration to GCL between days 1-3 (**Figure 1A, B**). BrdU^+^ cells were absent from PBS treated control group on day 1 and 3 (**Figure 1C**). The BrdU^+^ cells on day 1 at the INL co-expressed the Müller marker SOX9^17, 39^ (**Figures. 1D**, **Figure S4A**) confirming their Müller origin. On day 6, BrdU^+^ cells in GCL layer were RBPMS^+^ depicting their reprogramming to CiGN cells (**Figures. 1E, Figure S4B**). In summary, 6C induces *in vivo* Müller glia cell cycle entry, migration, and reprogramming into CiGN in 12-month-old DBA/2J mice.

**Figure 1.**
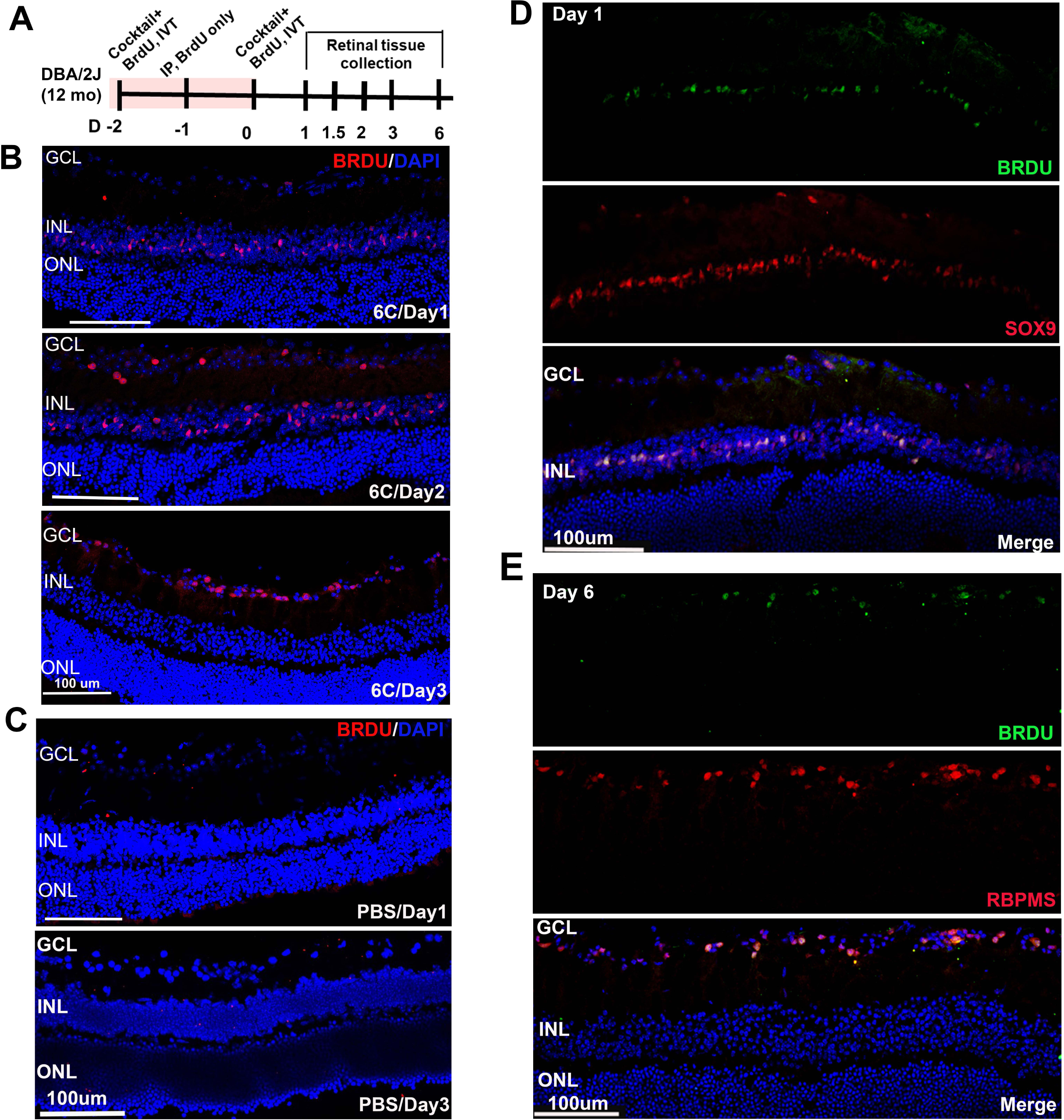
6C induces *in vivo* Müller glia cell cycle entry, migration to GCL and differentiation to CiGN cells in DBA/2J (12 mo old). **A.** Scheme for 6C injection and subsequent retinal tissue collection. **B.** BrdU^+^ cells generated in the INL on day 1 after 6C injection and subsequently moved to GCL on day 2 and 3. **C.** No BrdU^+^ cells were evident in PBS injected retina on day 1 and 3. **D.** BrdU^+^ cells in the INL are positive for Müller cell marker Sox9 on day1. **E.** migrated BrdU^+^ cells to GCL are positive for RBPMS on day 6.

### CiGN reprogramming proceeds through a SOX2-positive and SOX2-negative intermediate cellular state, like teleost fish

To investigate how 6C treatment induces reprogramming in DBA/2J mice, we analyzed SOX2 expression patterns during cellular transitions. In Medaka fish and Zebrafish, SOX2 plays a crucial role in the proliferation and neural commitment of Müller glia-derived progenitors upon injury^40^. Notably, SOX2 gain-of-function in the zebrafish retina can trigger a proliferative response even in the absence of injury^41, 42^. In mammals, including humans, SOX2 is crucial for maintaining postnatal Müller glia in a quiescent progenitor state and promoting neurogenesis in the central nervous system^43, 44^. Based on these insights, we hypothesized that SOX2 may play a role during 6C-induced reprogramming. To test this, we examined SOX2 expression at key time points—day 1 and day 1.5, when cellular transitions were observed in our BrdU injection experiment (**Figure 2**). On day 1, SOX2^+^ Müller glia in the INL were also BrdU^+^, indicating cell cycle entry (**Figure S5A**). Between days 1 and 2, we detected BrdU^+^SOX2^-^ cells (**Figure S5B;** blue arrowheads; **Figure S5C)** along with BrdU^+^SOX2^+^ cells (**Figure S5B;** yellow arrowheads; **Figure S5D**) in the INL and inner plexiform layer (IPL), suggesting that SOX2^+^ Müller glia may undergo asymmetric division, giving rise to BrdU^+^SOX2^-^ and BrdU^-^SOX2^+^ intermediate cells. We observed asymmetric nuclear divisions in the INL that generate BrdU^+^SOX2^-^ and BrdU^+^SOX2^+^ cells (**Figure S5E-G)**. We anticipate a subset of BrdU^-^ SOX2^+^ cells remain in the INL (possibly replenish the host Müller glia), while some BrdU^+^SOX2^+^ cells also migrated towards GCL. These results indicate that BrdU^+^ Müller cells proliferate in the INL, then migrate toward the GCL, transitioning through SOX2^+^ and SOX2^−^ intermediate cellular states like Zebra fish and Medaka respectively^40^.

**Figure 2.**
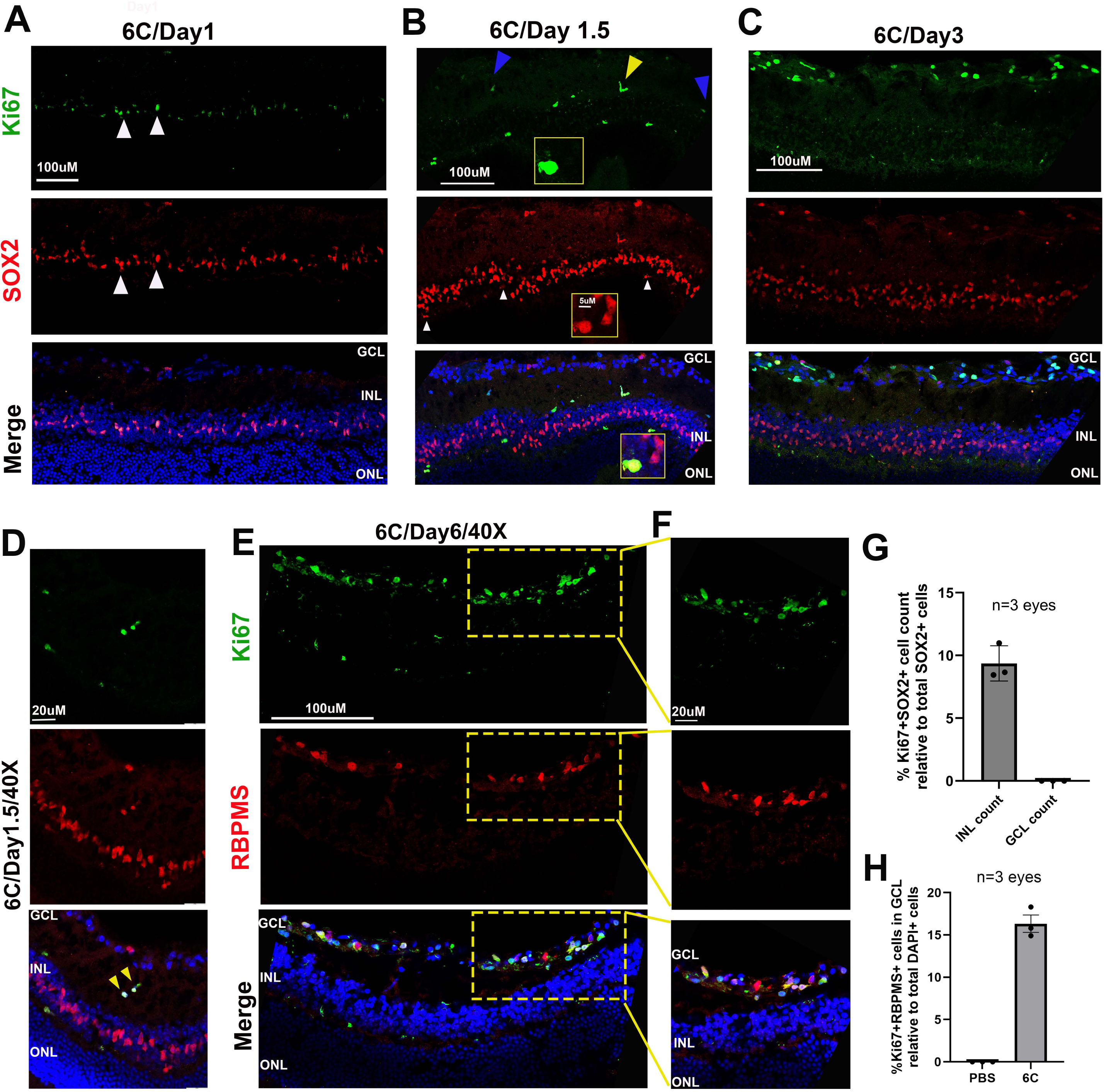
Induction of SOX2^+^ proliferative intermediate cells by 6C treatment, followed by their migration and differentiation into CiGN cells in 12-month-old DBA/2J mice. A-C. Generation and migration of Ki67^+^SOX2^+^ and Ki67^+^SOX2^-^ intermediate cells from the INL to the GCL between days 1 and 3. Inset: remaining SOX2^+^Ki67^+^ Muller cells in the INL between day1-2. **D.** High-magnification (40X) image showing cell migration of Ki67^+^SOX2^+^ intermediate cell through the IPL. **E, F.** Ki67^+^ cells co-expressing the RGC marker RBPMS on day 6. **G, H.** Quantification of Ki67^+^SOX2^+^ and Ki67^+^RBPMS^+^ cells in the INL and GCL following 6C induction (n= 3 eyes, 4 sections/eye, 3 fields/section).

To further validate that 6C treatment stimulates cell proliferation in the INL, we examined Ki67, a marker of actively dividing cells (excluding G0 phase), in the same retinas from days 1–2. On day1, we detected Ki67^+^SOX2^+^ cells in the INL (**Figure 2A)**, confirming their cell cycle entry. On day 1.5, we detected proliferating Ki67^+^SOX2^+^ (**Figure 2B**, yellow arrowhead, **2D, 2G)** and Ki67^+^SOX2^-^ (**Figure 2B**, blue arrowhead) in the IPL, consistent with their cell cycle entry and migration toward the GCL. At this stage we observed the remaining SOX2^+^Ki67^+^ proliferative Muller glia in the INL (**Figure 2B**, white arrowhead and insect). By day 3, these proliferating Ki67^+^ cells reached in the GCL (**Figure 2C, D**) and on day 6, these Ki67^+^ cells in GCL expressed RBPMS, confirming their CiGN identity (**Figure 2E, F-H**). Notably, a subset of Ki67^+^ cells in the IPL asymmetrically divide their nucleus and downregulated SOX2 expression during migration (**Figure S4C, D**). After reaching GCL on day 3, Ki67^+^ cells divided, lost their SOX2 and Ki67 expression, indicating their lineage commitment (**Figure 3A, C**, white arrowheads). On day 6, proliferating Ki67^+^ cells demonstrated lineage commitment, lost their proliferative capacity and exited the cell cycle, as evidenced by the presence of RBPMS^+^BrdU^+^Ki67^−^ cells in the GCL (**Figure 3D**, white arrowheads). Cell cycle exit indicated by BrdU^+^Ki67^-^ cellular stage similar to other published studies^45, 46^. Additionally, we observed migrated Ki67^+^SOX2^+^ and Ki67^+^SOX2^−^ intermediate cells in the optic nerve head region of 6C-treated retinas on day 3 (**Figure 3B**, red and white arrowheads). Altogether, these findings further confirm that CiGN regeneration proceeds through SOX2^+^ and SOX2^−^ intermediate stages derived from Müller cells, mirroring the process observed in teleost fish^40^.

**Figure 3.**
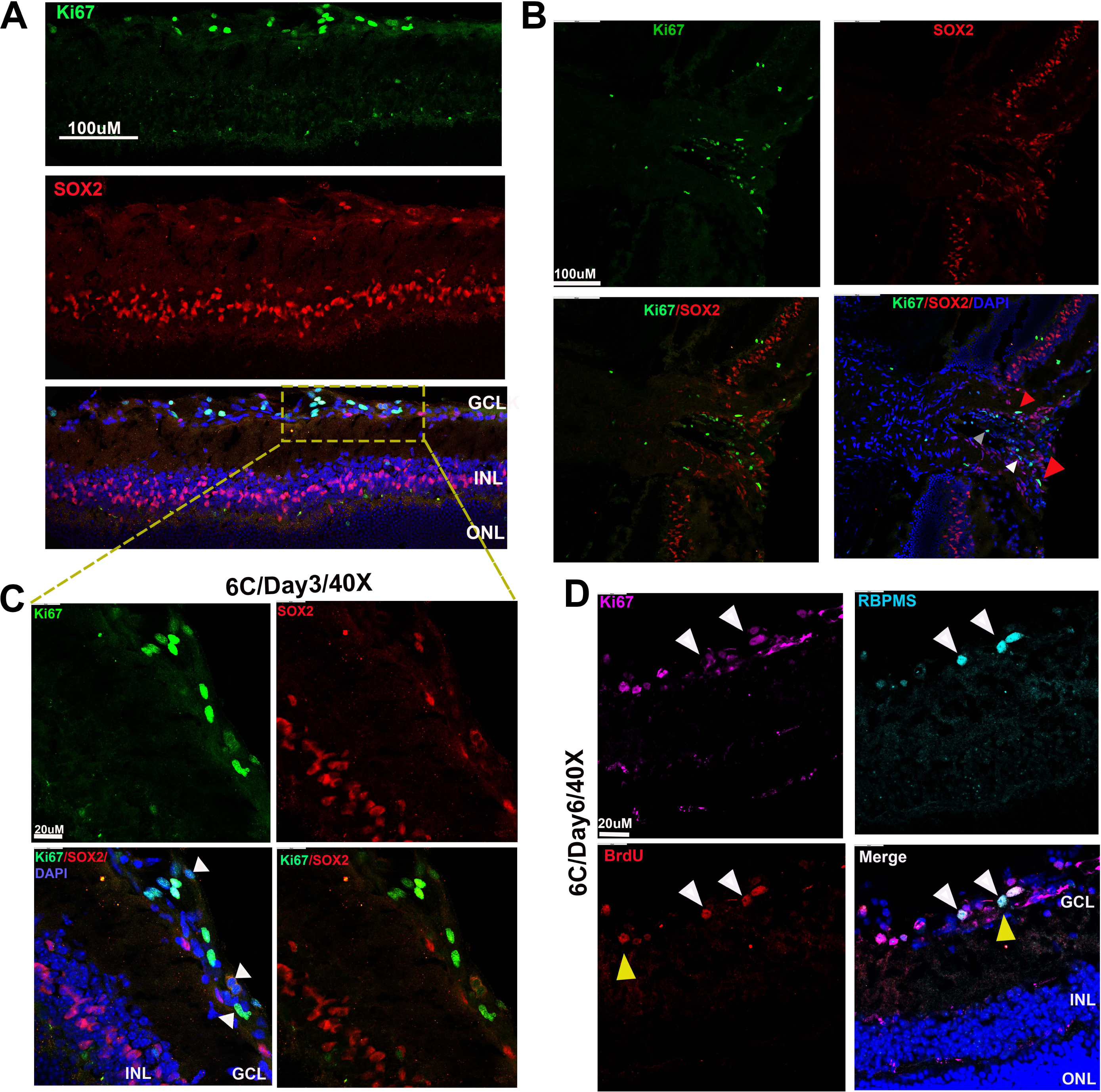
Loss of SOX2 expression in proliferative cells within the GCL and optic nerve head, followed by their differentiation into CiGN cells in DBA/2J mice after 6C treatment. **A.** on day 3, Ki67^+^ cells in the GCL lose SOX2 expression. **B.** High-magnification (40X) image showing reduced SOX2 expression (white arrowheads) in Ki67+ cells at GCL. **C.** Prospective migration of Ki67+SOX2+ (red arrowhead) and Ki67+SOX2-(white arrowhead) intermediate cells toward the optic nerve head region. **D.** Triple immunostaining showing the loss of proliferative capacity, cell cycle exit (possible G0 stage), and differentiation into CiGN cells in the GCL.

### 6C intravitreal administration rescued long-term visual functions post RGC injury

Next, we investigated if 6C mediated CiGN reprogramming could rescue visual functions after ocular hypertension induced RGC injury in 12-month-old DBA/2J mice using pattern ERG and VEP (pERG/pVEP). Baseline intraocular pressure (IOP) and pERG/pVEP were recorded in these mice from 7.5 to 11.5 months of age. We observed elevated IOP (**Figure S6A**) and a progressive decline in pERG/pVEP amplitudes between 7.5 and 11.5 months of age (**Figure S6B-E**). At age 12.5 months, we administered 6C or PBS intravitreally into these mice eyes (**Figure 4A**). By day 23 post-6C treatment, significant improvements in pERG/pVEP were observed. This pERG/pVEP recovery in DBA/2J mice lasting up to 4 months post 6C-treatment (**Figure 4B, C**). To investigate whether the observed rescue of visual function is injury-specific, we administered 6C after NMDA-induced RGC injury. NMDA (100 mM) was injected intravitreally, and 7 days later, mice were treated with either 6C or PBS. NMDA-injured mice exhibited a significant reduction in the number of RBPMS^+^ RGCs (**Figure S7A-D**) and reduced pERG/pVEP amplitudes on day 7 post-injection (**Figure 5A; 5B, C**, middle panel). We observed cellular events similar to DBA/2J in NMDA mice model after 6C treatment (**Figure S7E-H**). A notable improvement in pERG/pVEP amplitudes was observed post 6C administration, starting on day 16 and continuing through days 23, 30, 60, 90, 120, and 150 (**Figure 5B, C**, right bar graphs). By day 60, the pERG and pVEP amplitudes reached 63% and 89% of baseline levels, respectively, with improvements persisting through later time points. We noted no significant pERG/pVEP improvement until days 7 and 16 post 6C injections (**Figure 5B, C**, right bar graphs).

**Figure 4.**
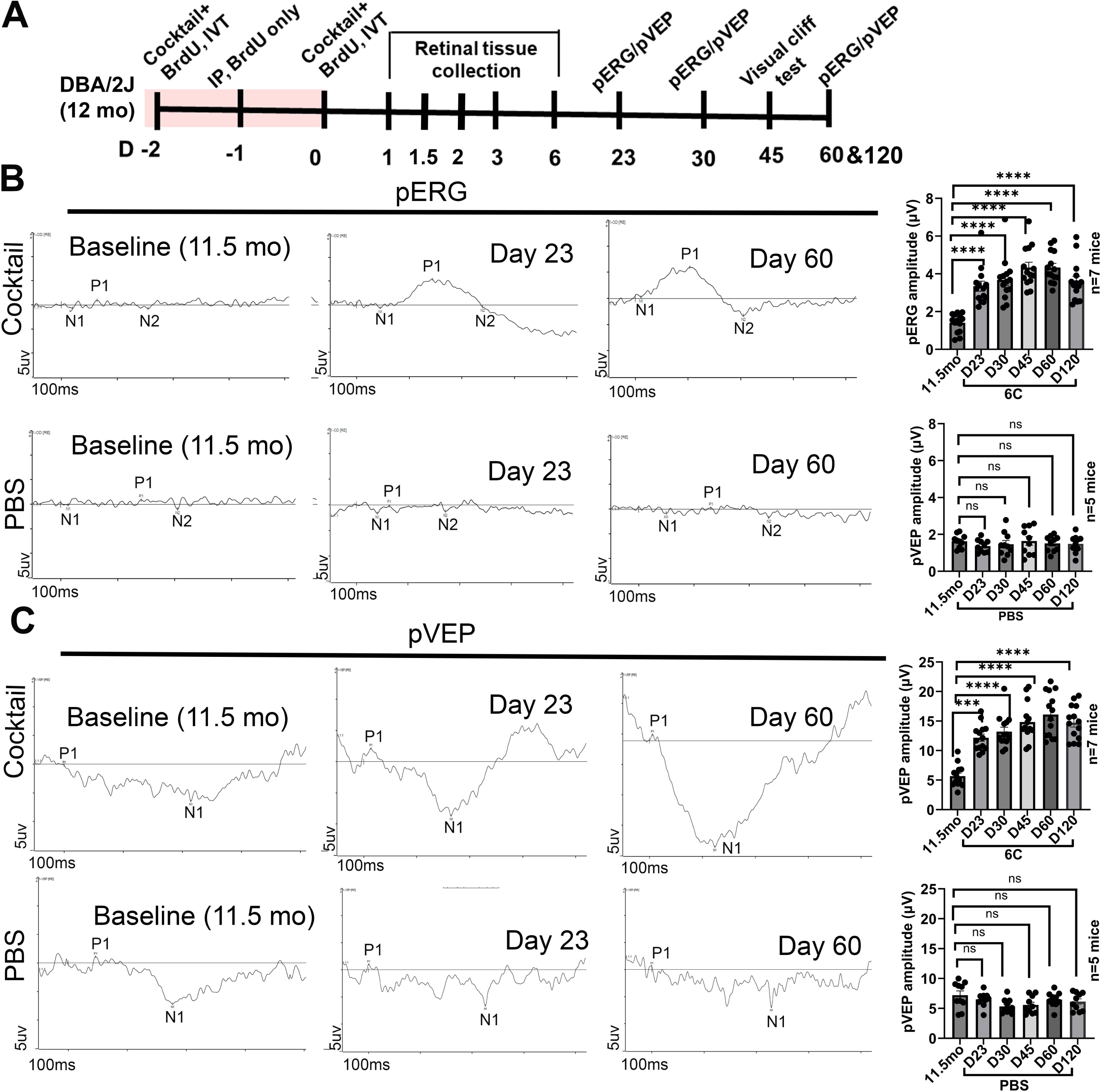
6C rescued long-term pERG/pVEP in 12-month-old DBA/2J mice. **A.** Timeline for functional measurements after 6C treatment in DBA/2J mice. **B.** Recovery of pERG upon 6C treatment. Upper panel: 6C treated. Lower panel: PBS treated. Representative pERG waveforms are from day 23 and day 60. **C.** pVEP measurements at the indicated time points. Upper panel: 6C treated. Lower panel: PBS treated. Representative pVEP waveforms from day 23 and day 60 are presented. All measurements were conducted longitudinally at indicated time points.

**Figure 5.**
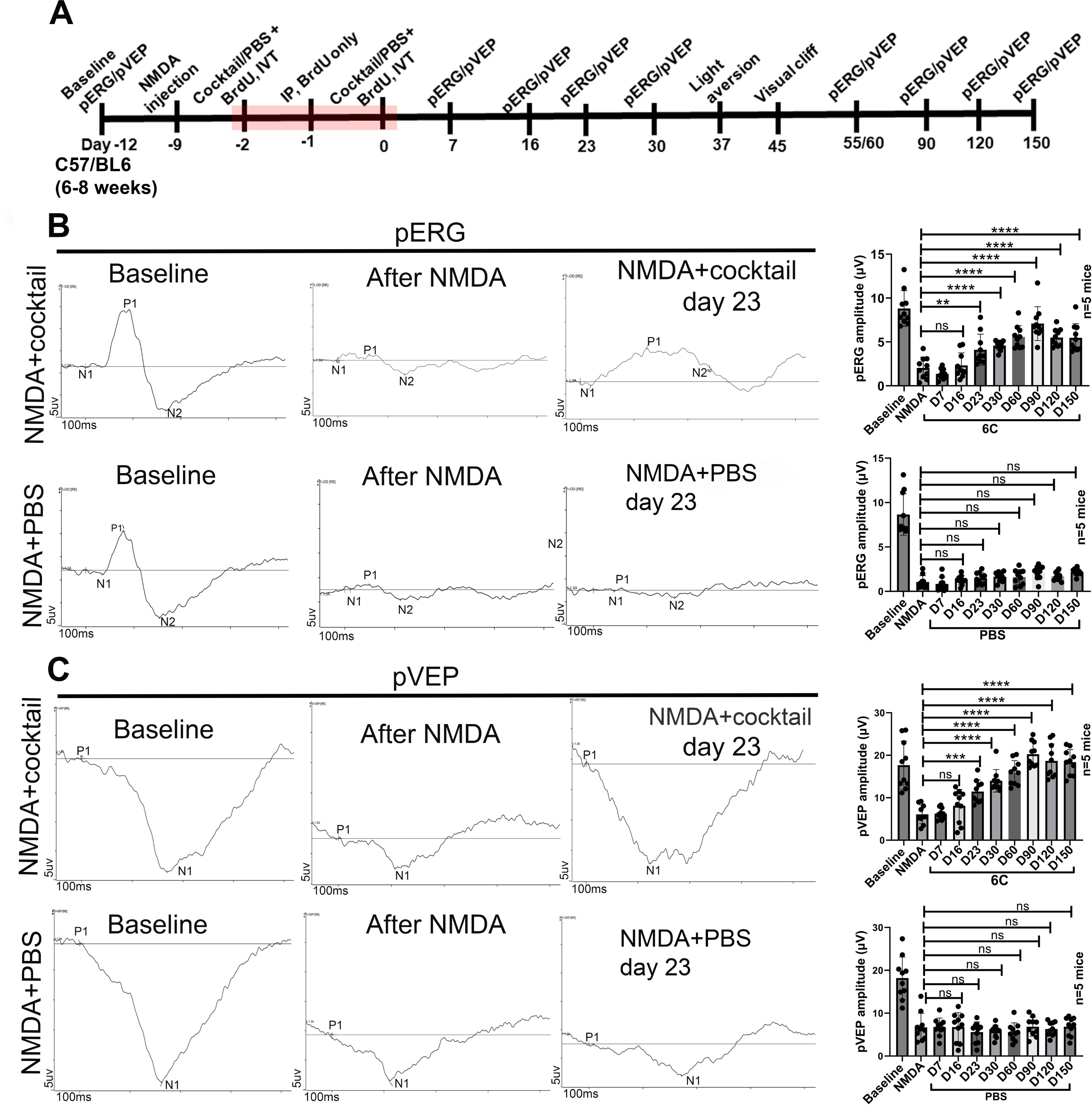
6C rescued long term pERG/pVEP after NMDA injury. **A.** Timeline for functional measurements after 6C treatment in NMDA injured mice. **B.** Recovery of pERG upon 6C treatment. Upper panel: 6C, lower panel: PBS. Representative pERG waveforms are from day 23. **C.** pVEP measurements at the indicated time points. Upper panel 6C, lower panel PBS. Representative pVEP waveforms from day 23 are presented. All measurements were conducted longitudinally at indicated time points.

Next, we conducted a longitudinal light avoidance test in the pERG/pVEP rescued NMDA injured animals following published methods to determine whether improved retinal function is sufficient to rescue visually evoked behaviour^47–49^. Normal mice exhibit a natural aversion to brightly illuminated areas, forming the basis for this test. All animals were placed in the apparatus for a duration of 300 seconds, and the proportion of time spent in the brightly lit (2800 lux, similar to cloudy outdoor environment) area was recorded. Wild type mice demonstrated a distinct preference for the dark compartment (194±14 sec), while PBS treated NMDA injured (123±12 sec) mice did not show such preference. The 6C treated mice exhibited a clear and statistically significant preference for the dark compartment (215±37 sec) as compared to PBS treated animals (**Figure S8A, B**). Additionally, wildtype (8±1.5 sec) and 6C treated groups (8±1.6 sec NMDA injured) showed similar latency to first entry to the dark chamber, while significantly higher latency time in PBS treated group in NMDA injured (38±5) (**Figure S8C**). Subsequently, we conducted longitudinal visual cliff test, a method employed to assess depth perception and the fear of crossing the deep side of a platform^50–52^. Typically, wild type mice exhibit an inherent tendency to avoid the deep side, favoring instead the shallow side of a visual cliff. Mice were positioned on the central platform between the deep and shallow sides, and their choices were meticulously documented. Our observations revealed that wild-type mice displayed a notable preference, with 96±3.9% choices to shallow side. The PBS treated NMDA injured mice exhibited a significantly diminished performance, with only 32±8.9% choosing the shallow side. Conversely, the 6C treated group demonstrated a more favorable outcome, with 72±10.9% of choices directed towards the shallow side in NMDA injured mice (**Figure S8D and E)**. In assessing the cumulative duration of stay on both the shallow and deep sides, it became evident that wild-type mice exhibited a distinct preference towards the shallow side. Conversely, the PBS treated NMDA injured mice showed no clear preference, indicating a compromised performance. Intriguingly, 6C-treated mice displayed a preference for the shallow side (**Figure S8F**). These findings strongly indicate a recovery of long-term visual functions in RGC injured mice after 6C mediated CiGN regeneration.

### Lineage tracing showed that CiGNs generated from Müller glia likely through asymmetric divisions and emerge during two different time windows

To lineage-trace CiGNs, we utilized NMDA-injured GLAST-CreERT2;ROSA26-flox-stop-flox (Ai9) reporter mice (**Figure S9A**) following the same reprogramming protocol as demonstrated in the NMDA and DBA/2J functional studies. The Müller-specific GLAST promoter-driven TdTomato expression (in nucleus and cell processes) within INL, along with the absence of TdTomato^+^ axons in the optic nerve, makes this model well-suited for lineage tracing and axon-targeting studies (**Figure S9B, C**). We did not observe TdTomato^+^ nuclear staining in the GCL of these mice. Upon 6C administration, Müller glia entered the cell cycle as early as day 1 post-injury, evidenced by BrdU^+^TdTomato^+^ cells in the INL, which subsequently migrated to the GCL by day 3 (**Figure S10A, B**). Next, we examined the presence of SOX2^+^ intermediate cellular stages in the NMDA model, as observed in our DBA/2J studies. On day 1.5, two distinct cell populations were identified in the INL: BrdU^+^SOX2^+^TdTomato^+^ (**Figure S10C**, white arrowheads) and BrdU^+^SOX2^+^TdTomato^-^ cells (**Figure S10C**, blue arrowheads). We observed that these cell types arose from asymmetric cytoplasmic divisions, similar to our findings in DBA/2J retinas (**Figure S10D**, yellow arrowhead). By day 2, BrdU^+^SOX2^+^TdTomato^+^ (**Figure S11A**, yellow arrowheads) and BrdU^+^SOX2^+^TdTomato^-^ **(Figure S11A**, blue arrowheads, cyan boxes) cells migrated through the IPL toward the GCL, with some already reaching GCL. At this stage, we observed additional asymmetric cytoplasmic divisions in the GCL, where BrdU^+^SOX2^+^TdTomato^-^ cells emerged from BrdU^+^SOX2^+^TdTomato^+^ intermediates (**Figure S11A**, purple arrowheads, yellow boxes). By day 6, both type of intermediate cells lost SOX2 expression in GCL and BrdU^+^TdTomato^-^ intermediates expressed RBPMS, indicating their potential differentiation into CiGNs (**Figure S11B, C**). However, at this stage, the migrated BrdU^+^TdTomato^+^ cells in GCL did not express RBPMS. By day 90, at the peak of pERG/pVEP functional recovery in NMDA functional experiment, a subset (but not all) of nuclear TdTomato^+^ CiGNs in the GCL began expressing RGC markers SNCG and RBPMS, suggesting their maturation into CiGNs (**Figure S12A-D, F, G**). We did not observe nuclear TdTomato expression in PBS-injected control eyes (**Figure S12E**). These results suggest Muller origin of CiGNs.

Next, we further validated the lineage tracing experiment using another transgenic mouse model in which the Müller cell specific Glast promoter drives expression of the SUN1 fusion protein in the inner nuclear membrane, enabling specific tracing of Müller glia fate following 6C administration after NMDA injury. Consistent with the Ai9 model, we observed BrdU⁺/GFP⁺ cells in the inner nuclear layer (INL) at day 1, indicating Müller glia cell cycle re-entry (**Figure S13A).** In contrast, PBS-injected controls showed no BrdU⁺/GFP⁺ cells in the INL at days 1 or 3 (**Figure S13B, C).** Between days 1.5 and 3, we detected migration of both BrdU⁺/GFP⁻ and BrdU⁺/GFP⁺ cells, suggesting that the BrdU-labeled cells appearing in the GCL originated from Müller glia (**Figure S14A-C**). By day 6, these BrdU⁺/GFP⁺ cells were distributed throughout the GCL (**Figure S14D).** Detailed analysis of retinas at day 1.5 post 6C injection revealed evidence of asymmetric nuclear division in the INL, likely giving rise to BrdU⁺ progenitor-like cells that undergo further division in the INL before migrating to the GCL (**Figure S15A, B** yellow arrowheads). At this stage we observed prospective BrdU+ dividing cells in the INL layer **(Figure S15A, B** white arrowheads). Together, these findings suggest that 6C treatment induces asymmetric cytoplasmic and nuclear division in Müller glia, generating BrdU⁺ progenitor-like cells that subsequently divide and migrate to the GCL.

### 6C induces axon regeneration in the optic nerve and establishes connections to the visual thalamus

Next, we examined if *in vivo* reprogrammed CiGNs in the GCL project their axons to the optic nerve in the same GLAST-Cre;Ai9 mice (post NMDA) 90 days after 6C treatment. We observed TdTomato^+^ axons passing through the optic nerve head and extended to the distal optic nerve in 6C treated eyes but not in PBS treated mice (**Figure 6A, B**). Further analysis of 6C treated eyes indicated expression of TUBB3 (a RGC axonal marker)^53^ in TdTomato^+^ axons that are passed the optic nerve via optic nerve head (**Figure 6C, D**)^54^. At this stage we also observed many TUBB3^+^TdTomato^-^ axons along with TUBB3^+^TdTomato^+^ axons in the optic nerve. We next carried out anterograde and retrograde tracing experiments to assess connectivity of TdTomato+ regenerated axons. Alexa 488–labeled Cholera Toxin B (CTB) was injected into the vitreous (anterograde) and LGN (retrograde) of 6C-and PBS-treated animals (**Figure S16A, B)**. In the anterograde tracing cohort, CTB transport was detected through TdTomato^+^ reprogrammed RGC axons at day 90 following 6C treatment (CTB was delivered intravitreally three days prior to sacrifice) (**Figure S16C-E**). In retrograde tracing, CTB^+^TdTomato^+^ axons were also detected in the optic nerve (**Figure S17**). Retinal analysis of retrograde-traced animals revealed three cell populations in the GCL: (a) CTB^+^ cells (green arrowhead), representing host RGCs; (b) a small number of TdTomato^+^ cells containing cytoplasmic CTB (yellow), suggestive of regenerated cells projecting to the LGN; and (c) TdTomato^+^ cells lacking CTB (red), indicating regenerated cells without LGN connectivity (**Figure S18A-C**). Additionally, we have performed retrograde tracing experiment in Glast-Cre;Sun1-GFP mice and analyzed the retina 4 months after 6C treatment (**Figure S19A, B**). We observed approximately 10% cells in the GCL containing both GFP and CTB staining indicating possible connection of GFP^+^ CiGN to LGN (**Figure S19C)**. Finally, we found TdTomato^+^ axon projections in dorsal lateral geniculate nucleus (dLGN) and ventral lateral geniculate nucleus (vLGN) in 6C treated mice brains but in PBS treated brains (**Figure S20A, B**). These results suggested that reprogrammed CiGNs extend axons to the optic nerve and establish connections with LGN.

**Figure 6.**
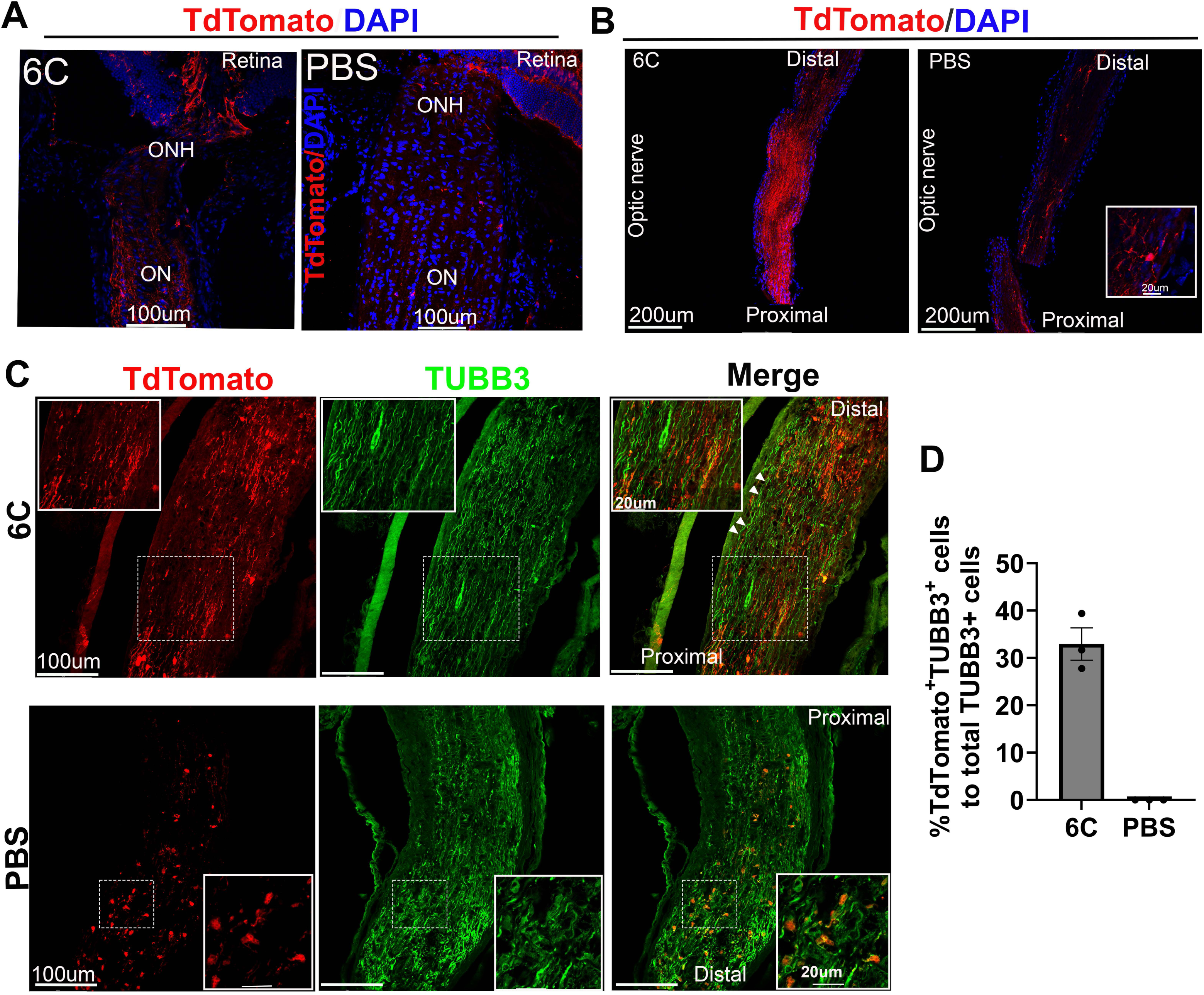
*In vivo* reprogrammed CiGN cells project axons toward optic nerve in GLAST-Cre;Rosa26-TdTomato reporter mice post NMDA injury. **A.** 6C vs PBS treated optic nerve head (ONH) showing TdTomato^+^ axons in the ONH projecting from reprogrammed CiGN cells. **B.** 6C vs PBS treated optic nerve showing TdTomato^+^ axon projections from proximal to distal direction. The PBS-treated nerve contains TdTomato+ cells with a star-shaped morphology, likely astrocytes (inset box). **C.** 6C vs PBS treated optic nerves showing TUBB3^+^TdTomato^+^ axons. Boxes showing enlarged TUBB3^+^TdTomato^+^ axons. **D.** Number of TUBB3^+^TdTomato^+^ axons passing optic nerve. n=3 eyes, 4 sections/eye, 3 fields/section were counted.

## DISCUSSION

Our study demonstrated the generation of photoreceptor-like cells using a defined chemical cocktail within 10 days^26^. Building upon this foundation, we explored additional molecules that could induce Müller glia reprogramming into retinal neurons. Among these, we investigated S24795 due to its role in neural progenitor cell survival and central nervous system neurogenesis^55^. Incorporating S24795 into the existing cocktail, we observed stimulated Müller glia cell cycle entry in INL, followed by migration to the GCL and differentiation into CiGN. Analysis of reprogramming intermediates in DBA/2J mice revealed that SOX2^+^ Müller cells proliferated in the INL within the first two days. A subset of these proliferating cells lost SOX2 expression either in the INL or the IPL before migrating to the GCL. Meanwhile, another subset of proliferating cells retained SOX2 expression during migration to GCL and eventually lost their SOX2 expression, This pattern of reprogramming resembles retinal regeneration observed in zebrafish and medaka fish, where SOX2^+^ and SOX2^-^ Müller-derived progenitors give rise to retinal neurons, respectively^40^. We anticipate that diminished SOX2 expression in the GCL restrict the regenerative capacity Müller-derived intermediate cells in the INL and GCL, while injury-associated signals likely guide their migration. We observed following differences in the mechanistic process of CiGN regeneration between the DBA/2J and NMDA models. (i) In the DBA/2J model, both SOX2-positive and SOX2-negative intermediate cells migrated to the GCL, whereas in the NMDA model, only SOX2-positive migratory intermediate cells were evident. (ii) We noted asymmetric nuclear divisions predominantly within the INL and some in IPL in the DBA/2J model, while in the NMDA-injured retina, asymmetric divisions occurred in both the INL and GCL. (iii) We observed both BrdU^+^SOX2^+^ (S-phase positive) and Ki67^+^SOX2^+^ (all cell cycle stage positive except G0) cells in the INL and IPL of the DBA/2J mice. However, we observed only BrdU^+^SOX2^+^ but not Ki67^+^SOX2^+^ cells in the NMDA model. This variation in regenerative response may be attributed to NMDA-induced modifications of Müller cell proteome through methylene amine-mediated proteomic alterations, as reported in previous studies^56, 57^ or injury context in DBA/2J retina may influence regeneration upon 6C treatment^58^. We did not observe the expression of BRN3A or BRN3B during the reprogramming process at any time points, indicating that the CiGNs may arise via a pathway that bypasses these conventional RGC transcription factors through an alternative reprogramming route in line with a previous finding^59^.

While CiGN reprogramming was evident on day 6, functional recovery measured by pERG/pVEP began on day 23 post-6C injection. This delay may be attributed to the limited number of CiGN axons passing through the optic nerve head on day 6 (**Figure 6A**), requiring additional time for newly generated CiGN axons to establish connections with brain targets through the distal optic nerve. The presence of both TUBB3^+^/TdTomato^-^ axons along with regenerated TUBB3^+^/TdTomato^+^ axons in the optic nerve (**Figure 6C**), coupled with a relatively greater improvement in pERG (>60%) compared to the modest CiGN reprogramming (>15%), suggests that mechanisms beyond endogenous regeneration may contribute to vision rescue which requires further investigation. We anticipate following mechanisms based on our results: (i) Small molecules may pass into the optic nerve through glymphatic pathway which induces the reprogramming of optic nerve glia into bridge neurons and may establish a neuronal relay mechanism that connects with CiGN axons for sending light signals to the brain^60, 61^ (**Figures S17B and S22)**. (ii) Host RGCs may convert to a partially reprogrammed or a youthful state that potentially facilitate TUBB3^+^/TdTomato^-^ host RGC axon regeneration and vision recovery similar other studies^62–65^ (**Figure S21**). Increased number of SOX2^+^RBPMS^+^ cells in the GCL upon 6C treatment (**Figure S21A-C**) and induction of youthful pathways in intermediate clusters during *in vitro* CiRGC reprogramming supports this hypothesis (**Figure S21D**,)^66^. Importantly, the absence of pERG improvement on early time points (days 7, 16), followed by gradual increase, thereafter, suggests that neuroprotection is an unlikely mechanism underlying the observed visual recovery (**Figure 5B, C;** right bar graphs). Moreover, 6C induced material transfer between Muller cell endfeet and host RGCs may be a mechanism for generation of abundant TdTomato+ RGC axons along with CiGN axons in the optic nerve. Altogether these findings suggest a complex neuron-glia interplay within the optic nerve and associated brain regions may be involved in functional outcomes, warranting further investigation.

In summary, this study introduces a chemical-only approach that effectively triggers *in vivo* CiGN reprogramming through SOX2^+^ and SOX2^-^ intermediate stages and restores vision following RGC damage caused by ocular hypertension and NMDA-induced injury. This method presents a promising alternative to cell transplantation, offering a potential therapeutic strategy for optic neuropathies such as glaucoma where RGC loss is the final common endpoint.

## MATERIALS AND METHODS

### Animal models and lineage tracing

All animal studies and animal care were performed in accordance with relevant guidelines and regulations approved by IACUC at Children’s Hospital Los Angeles. Preliminary experiments were conducted on C57BL/6J mice (Jackson Lab.). For linage tracing experiments Glast-creERT2/ROSA26 mice line generated by crossing i) Tg(Slc1a3-cre/ERT)1Nat/J (stock # 012586) ii) B6.Cg-Gt(ROSA)26Sortm9(CAG-tdTomato)Hze/J (stock # 007909, Ai9). Both male and female mice were used in all experiments with equal frequency. All the *in vivo* experiments were conducted on 6-8 weeks old mice.

### DBA/2J mice glaucoma model

Three months old DBA/2J mice purchased from Jackson lab (stock # 000671) and kept in animal care facility at Children’s Hospital Los Angeles. Intraocular pressure was measured with Tonolab rebound tonometer (iCare, USA) at 3 months and 11.5 months of age (**Figure S7A**). Progressive decline in pERG/pVEP recorded at 7.5, 8.5, 10.5 and 11.5 months of age. Maximum decline in pERG/pVEP recorded at 11.5 months (**Figure S7**). Formal experiment started 12 months of age.

### Generation of RGCs injury mice model and intravitreal injection of small molecules

All intravitreal injections were performed under isoflurane anesthesia at CHLA animal care facility. For mice NMDA model, NMDA powder (Sigma-Aldrich) was dissolved in PBS to prepare 100 mM solution and injected under a dissection microscope placed inside a BSL-II laminar flow. Before the injection, 1% tropicamide was applied to dilate pupils and injections were carried out with a 10 µl Hamilton syringe mounted with a 30-gauge needle (Small Hub RN Needle, Hamilton). 1.5 µl of NMDA solution injected in intravitreal space at an angle of 45 degree with the lens. Small molecule cocktail containing V (10 mM), C (96 μM), R (40 μM) and F (200 μM) I (200 μM) supplemented with Noggin (800 ng/mL) IGF-1 (8 µg/mL) S24795 (1 mM) and BrdU (1 mg/mL) in 50 µl PBS were prepared and 1.5µl/eye was injected according to the scheme described in result section (Figure 5 A and 6A). One additional BrdU IP injection was performed 4-6h before the tissue collection. Same concentrations were used for DBA/2J model.

### Generation of chemically induced Retinal ganglion cells *in vitro*

Human primary Muller cells (HMC, Creative Biolabs, Cat# NCL2110P081) were seeded in 24 well plates (0.1 % gelatin coating) and cultured overnight. Next day, the normal growth medium was replaced with Retinal differentiation medium (RDM1, containing small molecules V (0.5 mM), C (4.8 μM), R (2 μM), F (10 μM). On day 2, ISX9 (Is) (20 µM) together with VCRF (same concentration) added to cells in RDM2 medium. On day 3, Taurine (T), Retinoic acid (R) and sonic hedgehog (S) also added together with other small molecules in RDM2 medium. Cells were collected for molecular assays, sequencing studies and immunofluorescence staining on day 4.

### Real time qPCR

Total RNA was extracted and treated with DNAase-I according to the manufacturer’s instructions (Zymo Research, Cat# R1050). Up to 2 μg of RNA was reverse transcribed to cDNA using a High-Capacity cDNA Reverse Transcription kit (Applied Biosystems, 4368814) in BIO-RAD thermal cycler (T100). The qPCR was performed using Fast SYBR Green Master Mix (Applied Biosystems, 4385612) in Quant studio 7 Real Time PCR machine. The fold change compared to starting cells was calculated by 2 -ΔΔCT method after normalizing with glyceraldehyde 3-phosphate dehydrogenase.

### Immunostaining of tissue sections and cells

Enucleation was performed after CO2-mediated euthanasia. Enucleated eyes were fixed for 15 min in 4% paraformaldehyde (PFA) in phosphate-buffered saline (PBS). After fixation, the cornea was removed, and eye cups were sequentially incubated in 15% sucrose and then 30% sucrose for overnight at 4°C. Eye-cups were then frozen in optimal cutting temperature compound (O.T.C.) and 30 μm cryosections were prepared. For immunostaining, sections were washed two times with PBS, three times with 0.1% PBST and permeabilized with 0.25% PBST for 15 minutes. After blocking for one hour, sections were incubated with primary antibodies overnight at 4°C. Next morning, sections were washed five times with 0.1% PBST and incubated with secondary antibodies (Alexa 488 or 594) for 90 minutes. Lastly, slides were washed 10 times with 0.1% PBST and mounted in Fluoromount-G containing DAPI and dried for two hours.

For Immunofluorescent staining of reprogrammed RGCs, cells were fixed for 20-25 min in 4% paraformaldehyde (PFA) in phosphate-buffered saline (PBS) and Permeabilized for 30 mints in 0.25% PBST. After blocking for one hour at room temp., cells were incubated with primary antibodies overnight at 4°C. Next morning, cells were washed five times with 0.1% PBST and incubated with secondary antibodies for 90 minutes. The cells were then washed with 0.1% PBST and incubated with DAPI solution (1ug/ml). Images were captured on Leica DMi8 confocal microscope at Imaging core facility, CHLA. Cell counting was performed using ImageJ and Adobe photoshop. For regenerated axon count, every fifth section was counted, and 4 sections were included per eye (n=3 eyes). For BrdU immunostaining, antigen retrieval was performed following permeabilization. After incubation with 0.25% PBST for 15 minutes, the tissue sections were sequentially treated with 1N HCl, 2N HCl, and phosphate/citric acid buffer (pH 7.4) for 10 minutes each. Sections were then washed three times with 0.1% PBST, blocked for 2 hours at room temperature, and incubated overnight at 4°C with the anti-BrdU primary antibody. After three additional washes with 0.1% PBST, sections were incubated with a fluorescently labeled secondary antibody for 90 minutes at room temperature. Finally, the slides were washed 10 times with 0.1% PBST, mounted in Fluoromount-G containing DAPI, and air-dried for 2 hours before imaging

### Transcardiac Perfusion, Brain and optic nerve Collection

ketamine/xylazine mixture (80 mg/kg ketamine, 10 mg/kg xylazine) was administered via intraperitoneal injection. Once the animal reached a surgical plane of anesthesia, a lateral incision was made just beneath the rib cage. Additional incisions were made from the xiphoid process along the ventral ribcage, and the thoracic field was exposed. The ribcage was reflected to expose the heart, and the pericardial sac was opened. The heart was secured, and a 25G syringe needle was inserted into the left ventricle. The right atrium was cut, and perfusion with PBS was initiated, followed by 4% PFA. The mouse was decapitated, the skull was opened, and the brain was removed and placed in 4% PFA overnight at 4°C. Optic nerves were collected after transcardiac perfusion and fixed in ice-cold 4% PFA for 20-30 minutes.

### Single cell RNA Sequencing

The feature-barcode matrices were obtained by aligning reads to the human genome using CellRanger v7.1.0. Raw feature barcode matrix files were run in a coding free open-source automated software Cellenics (Bioimage) according to manufacturer’s instructions [Cellenics User Guide (biomage.net)]. First, we processed the project to convert the count matrices into a Seurat object. Data processing pipeline will be triggered automatically after the successful generation of a Seurat object. The automated data processing pipeline values are established according to the current best practice and spread of each sample. This data processing module contains classifier filter, cell size distribution filter, mitochondrial content filter, number of genes vs UMI filters, doublet filter, data integration and configure embedding. After successful data processing, UMAPS (gene expression) were generated using the functions categorical embedding and continuous embedding. Heatmap and violin plot function in the software were used to generate these plots. Additionally, we have performed trajectory analysis for CiGN cells using the in-built monocle3 based function in the software. We first calculated the root nodes using this function and then selected the root nodes inside our cell cluster of choice (anticipated progenitor population) and then ran the function. For multiple scRNA-Seq data comparison, we first uploaded cell ranger derived raw feature barcode matrix files for all samples and then processed the data simultaneously using automatic processing functions. To generate cell POU4F1 or POU4F1^+^RBPMS^+^ cell sets data exploration function was used, and custom cell sets were generated after selecting gene or genes of interests. Cell type annotation was performed using an automatic function of the program after selecting human eye as reference tissue type.

### Light aversion behavior test

The light/dark apparatus consists of black opaque (100%) acrylic test chambers (30.48 × 15.24 × 30.48 cm (length, width, height)). This chamber was further divided into equal-sized compartments (15.24 × 15.24 × 30.48 cm) by the addition of an insert, to create a dividing wall in the center. Further, to create light and dark zones, one compartment was illuminated with light (2800 lux, similar to cloudy outdoor environment) and the other compartment was kept dark (∼ 0.1 lx). The light and dark compartments were connected by an opening (5 × 5 cm). The position of the mouse within the apparatus was recorded using a camera. Mice were maintained in the testing room for 2 hrs in ambient light (200 Lux) conditions in their home cage with free access to food and water. Each mouse was allowed to habituate to the testing apparatus (both sides) for 10 min while in ambient light. After habituation, one side of the apparatus was illuminated with light at around 2800 lux, and the mouse was allowed to roam freely between each compartment for 5 min.

### Visual cliff test

The apparatus was purchased from Conduct Science, Boston, MA, comprised a transparent acrylic chamber measuring 62 cm x 62 cm x 19 cm, featuring a checkered pattern insert. This chamber is divided by a central platform, 1.5 inches high and 2 inches wide, creating two sections. One side is shallow, with a checkered pattern immediately beneath the platform, while the other side is deep, with the same checkered pattern positioned approx. 2 feet below, producing a visual depth illusion. Both sides of the setup are evenly illuminated with a light intensity of approximately 2000-2500 lux. For shallow or deep choice experiments, mice were placed on the central platform, and their movements were captured using a video camera. Each mouse undergoes five trials.

### Pattern ERG and pattern VEP

For pERG and pVEP, anesthetized animals were placed on a feedback control heated stage (37°C) and were recorded by selecting and running specific protocol using default setup (Diagnosys LLC, Boston, MA). 1 % tropicamide eye drops were used to the pupil and GenTeal tear eye drops were used to prevent corneal desiccation. 6-8 weeks C57/BL6 mice (Jackson lab) were used for baseline measurement. Animals with stable baseline were used for NMDA injections (100 mM, 1.5μl/eye). Non-injected wild type mice were used as control. Eyes showed reduction in both pERG and pVEP amplitude after NMDA injection, were used for chemical cocktail injection following the protocol described in Figure 3a. Briefly, under red light pERG responses were recorded by touching pattern stimulator to cornea of one eye and a full filed stimulator as a fellow eye reference electrode according to manufacturer’s instructions (https://www.diagnosysllc.com/wp-content/uploads/2021/03/Celeris-Pattern-Stimulator-Application-Note-d.pdf). After pERG same mice were used for pVEP measurement by changing the electrodes in appropriate channels according to manufacturer’s instructions. For pVEP, responses were recorded by touching pattern stimulator to the cornea, a needle electrode placed sub dermally in mice snout and the reference electrode to the back of the head Then a pVEP protocol was ran using default parameters. Independent pERG or pVEP responses were obtained by default averaging system. Data (P1 and N1 amplitude values) were exported from the machine using Espion V6 software (Diagnosys, LLC). Webforms were exported as graphs.

### Retrograde and anterograde tracing with CTB

Retrograde tracing was performed by injecting cholera toxin subunit B (CTB; Thermo Fisher, Cat# C22841) into the lateral geniculate nucleus (LGN) of mice. Animals were anesthetized with isoflurane or a ketamine (100 mg/kg)/xylazine (10 mg/kg) mixture and positioned in a stereotaxic frame. Following scalp shaving and disinfection, a midline incision was made to expose the skull, which was cleaned with 3% hydrogen peroxide. Using bregma as the reference, stereotactic coordinates for the LGN were identified (approximately 2.3 mm anterior–posterior, ±2.1 mm medio-lateral, and 2.8 mm dorso-ventral). A small cranial opening was drilled, and 100–300 nL of 1% CTB solution was injected using a Hamilton syringe at a rate of 10–50 nL/min. The needle was left in place for 2–5 minutes to minimize backflow before withdrawal. The incision was closed with surgical adhesive or sutures, and mice were allowed to recover on a heating pad with appropriate postoperative monitoring and analgesic care. For anterograde tracing, 1–1.5 µL of 1% CTB was injected into the vitreous of the eye using a 32G Hamilton micro syringe under a dissection microscope. Animals were maintained for 72 hours and 2 weeks post-injection to allow tracer transport before euthanasia and tissue collection for anterograde and retrograde transport respectively.

### Statistical analysis and fluorescence quantification

The data are expressed as mean ± standard error of the mean (SEM). Statistical significance was assessed through Student’s t-test and one-way or two-way ANOVA using GraphPad Prism Software (version9), with corresponding P value summaries shown in the figures. For fluorescence quantification, images were analyzed using ImageJ.

## DATA AVAILIBILITY STSTEMENT

All data that supports the findings of this study are available from the corresponding author upon reasonable request. All sequencing data described in this work have already been deposited in the Gene Expression Omnibus (scRNA-Seq: GSE253807) and will be made public upon the publication of this article.

## ACKOWLEDGEMENTS

B.M. acknowledges grant support from following agencies: Knights Templar Eye Research Foundation, Glaucoma Research Foundation, Children’s Hospital Los Angeles, California Institute for Regenerative Medicine, CHLA office of the Technology Transfer, The Leon and Carol Ellison Research Career Development Award. We thank the stem cell analytics core, cellular imaging core, special biology and genomics core at Children’s Hospital Los Angels, NEI funded P30 core at USC. We thank Drs. David Cobrinik, Thomas Lee, Aaron Nagiel, Mark Borchert and Jesse Berry for critical review of the results.

## AUTHOR CONTRIBUTIONS

B.M. designed experiments; B.M, wrote the manuscript; R.M.S.; S. M.; A, S.; R. L., S. H.; P.T.; and BM: performed experiments. B.M.; R.M.S.; analyzed data.

## DECLEARATION OF INTERESTS

All other authors declare no competing interest. B.M. and R.M.S. are listed as inventors in a provisional patent application submitted to the office of technology and commercialization at Children’s Hospital Los Angeles.

## Figure Legends

**Figure S1.**
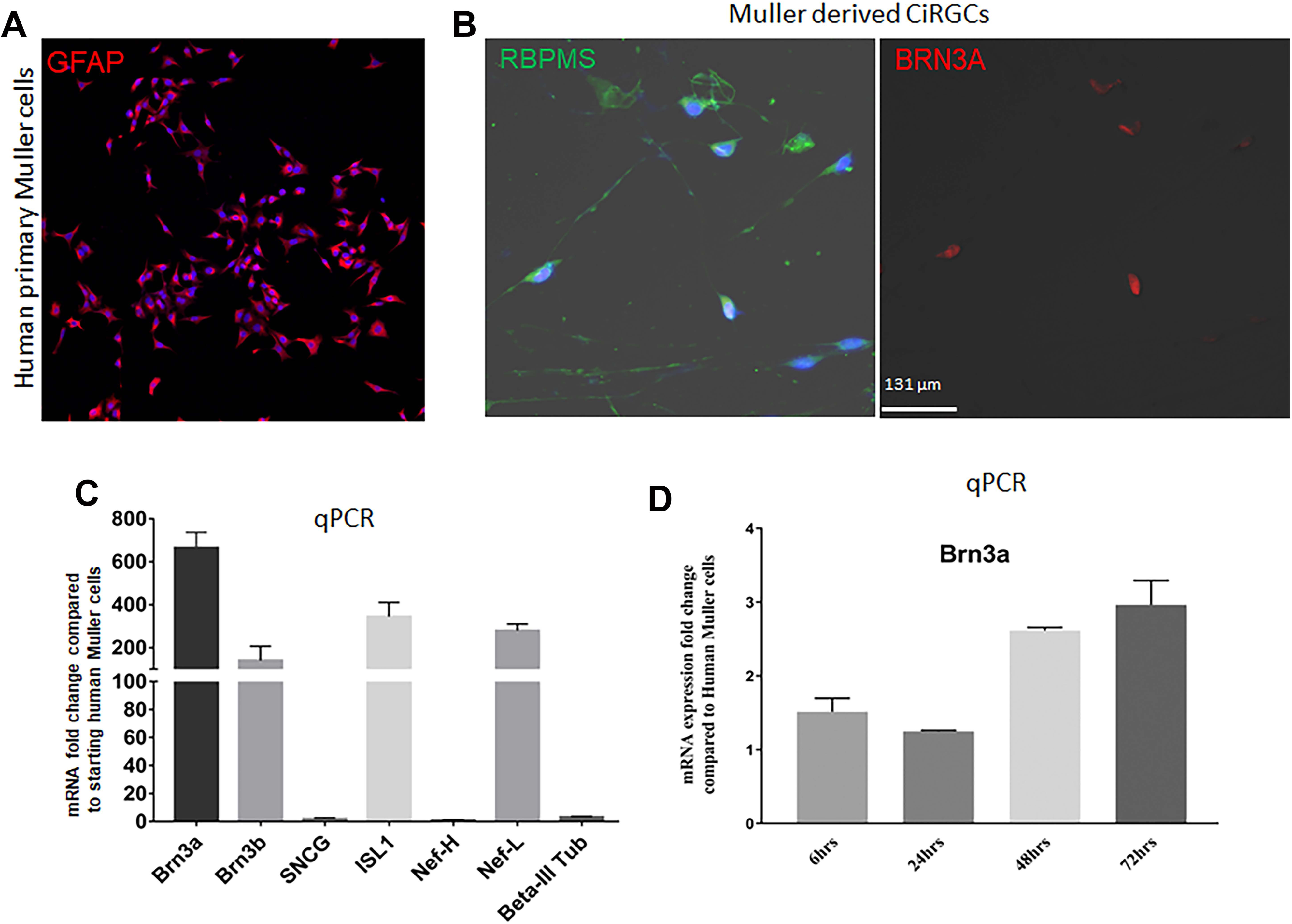
*In vitro* generation of CiRGCs from human Müller cells. **A.** Expression of GFAP in human primary Müller cells. **B.** RBPMS (green) and BRN3A (red) staining in Md-CiGN cells 4 days after chemical cocktail treatment. **C.** qPCR showing increased expression of major RGC specifying genes in CiGN cells on day 4 (Ct values less than 30 were considered for the analysis for all samples). **D.** Time course expression measurement of *Brn3a* mRNA by qPCR in CiGN cells.

**Figure S2.**
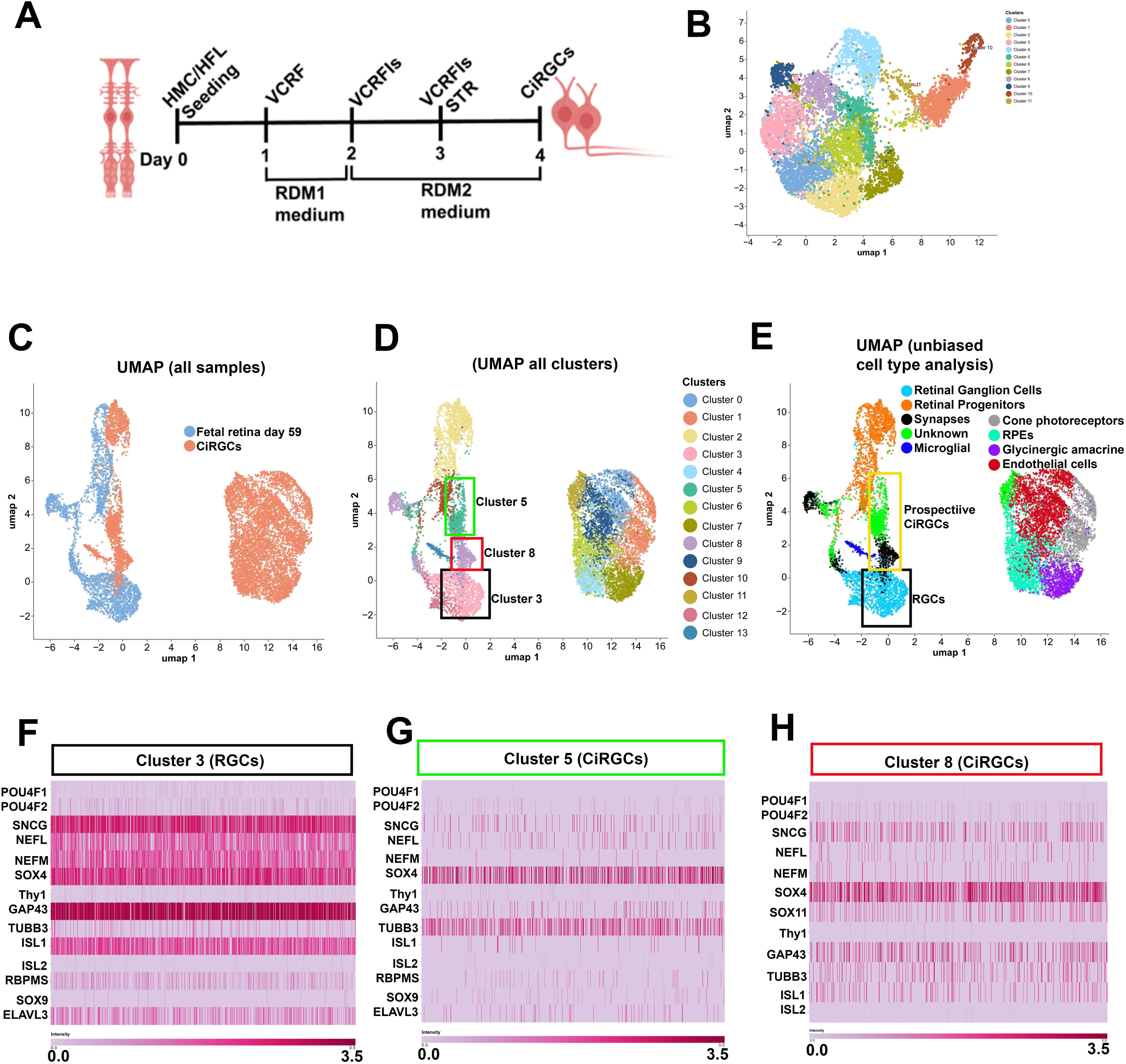
Comparison between *in vitro* CiRGCs and native RGCs from fetal retina d59 (scRNA-Seq). **A.** Scheme showing CiGN generation from human primary Müller cells (HMC). **B.** UMAP analysis after CiGN total cell population. **C, D.** UMAP analysis of CiGN cells total cell population and fetal retina day 59. A total of 13 clusters were found after embedding. **E.** Unbiased cluster annotation showing various retinal cell types and an unknown population (prospective CiGN cells) in chemically induced population. CiGN population fall near the native RGCs from fetal retina. **F.** Heatmap for RGC specific gene expression in clusters 3 (native RGCs) found in fetal retina day 59. **G, H.** Heatmap for RGC specific gene expression in clusters 5 and 8 (CiGN cells).

**Figure S3.**
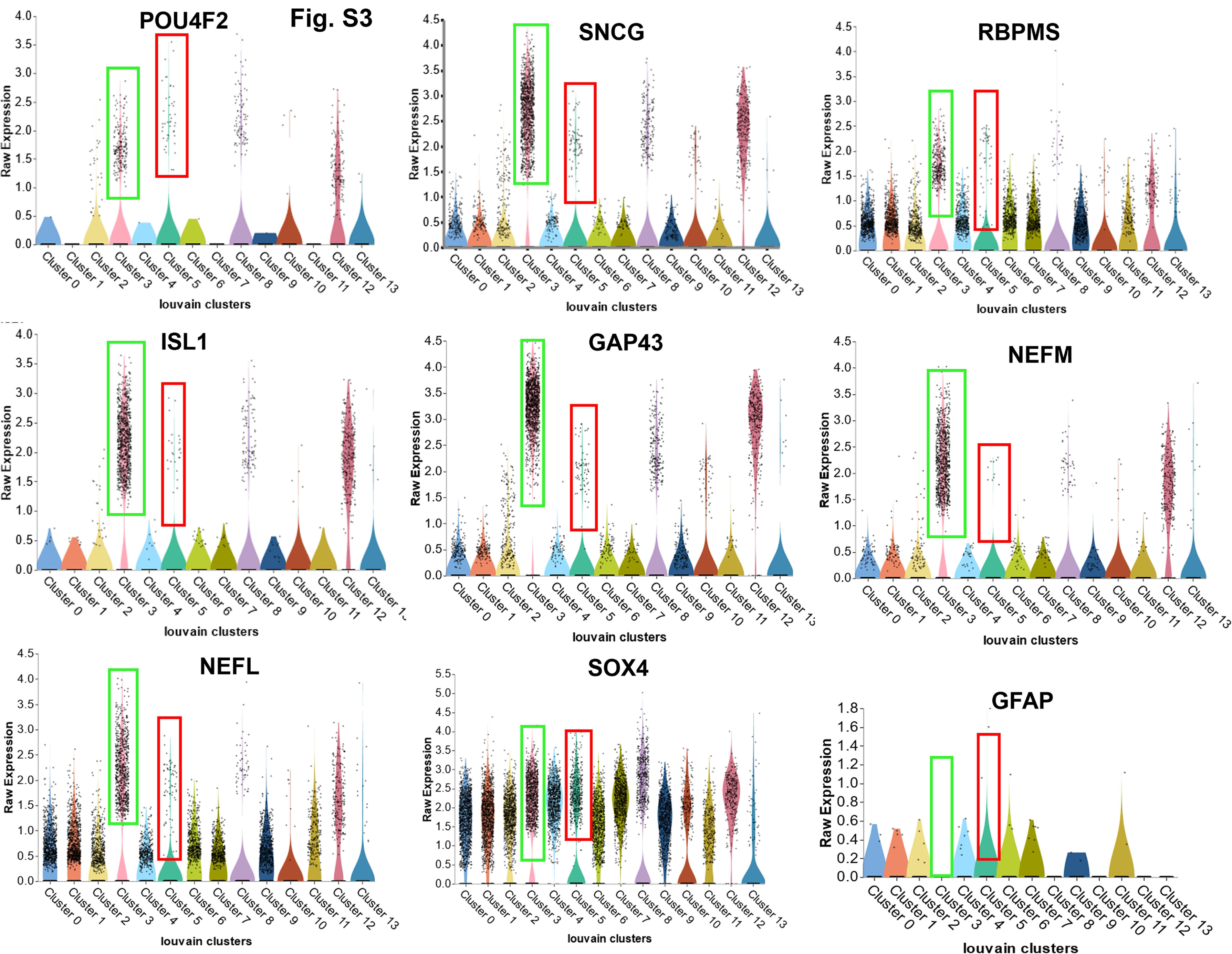
Comparison of RGC specific gene expression (violin plot) levels among different clusters after embedding *in vitro* CiRGCs total cell population and fetal retina d59 data sets. Green box: native RGCs; solid red boxes: CiRGCs

**Figure S4.**
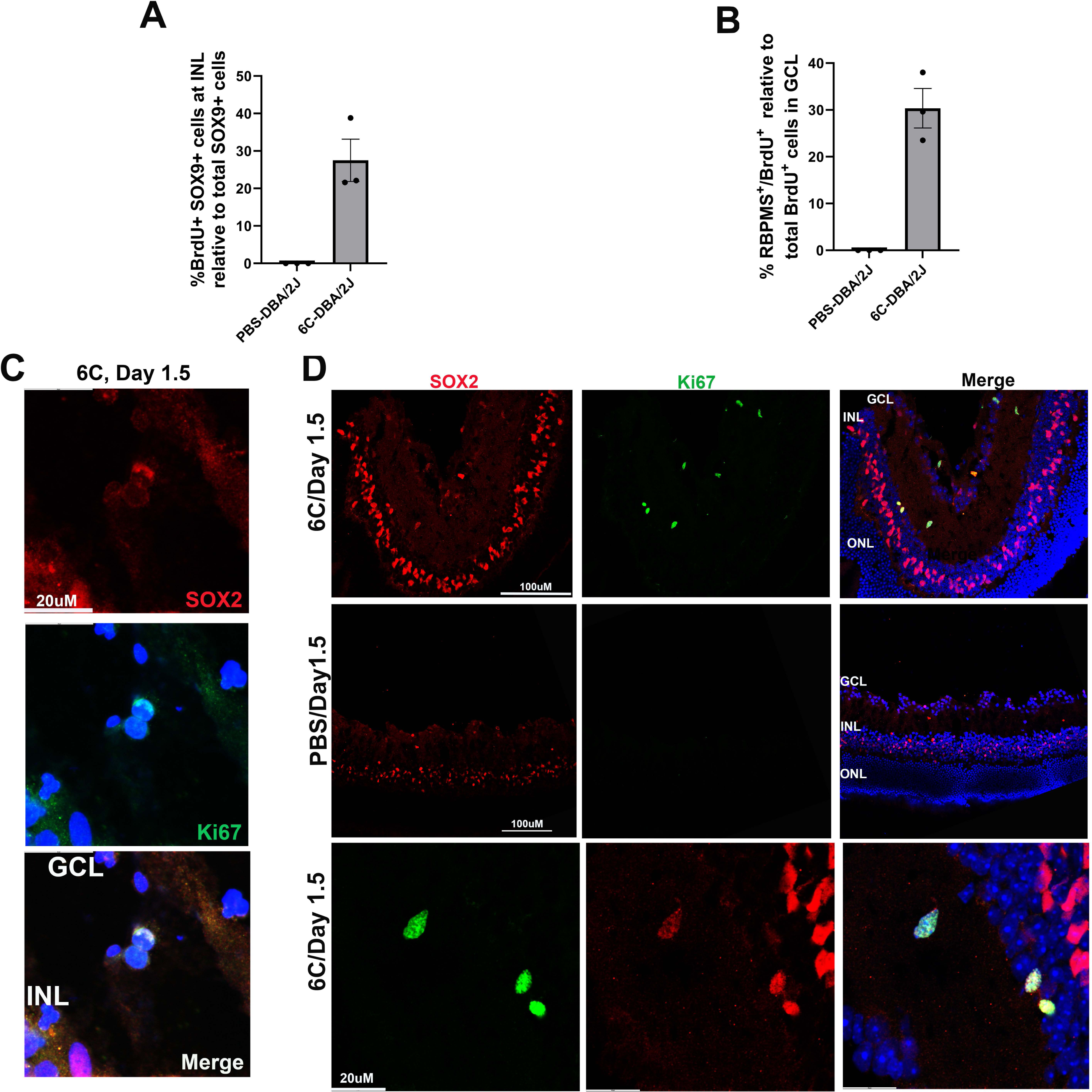
Quantification of *in vivo* reprogrammed cells and SOX2 expression dynamics in proliferating cells following 6C induction in DBA/2J mice. **A.** Percentage of BrdU+SOX9+ in the INL on day1 after NMDA injury and DBA/2J model. n=3 eyes, 5 sections/eye, 3 fields/section were counted. **B.** Percentage of BrdU^+^RBPMS^+^ cells in the GCL on day 3 after NMDA injury and DBA/2J model. n=3 eyes, 5 sections/eye, 3 fields/section were counted. **C.** Image for SOX2+ cell division during migration through IPL on day 1-2. **D.** A subset of Ki67+ proliferative cells gradually lose SOX2 expression as they migrate through the IPL.

**Figure S5.**
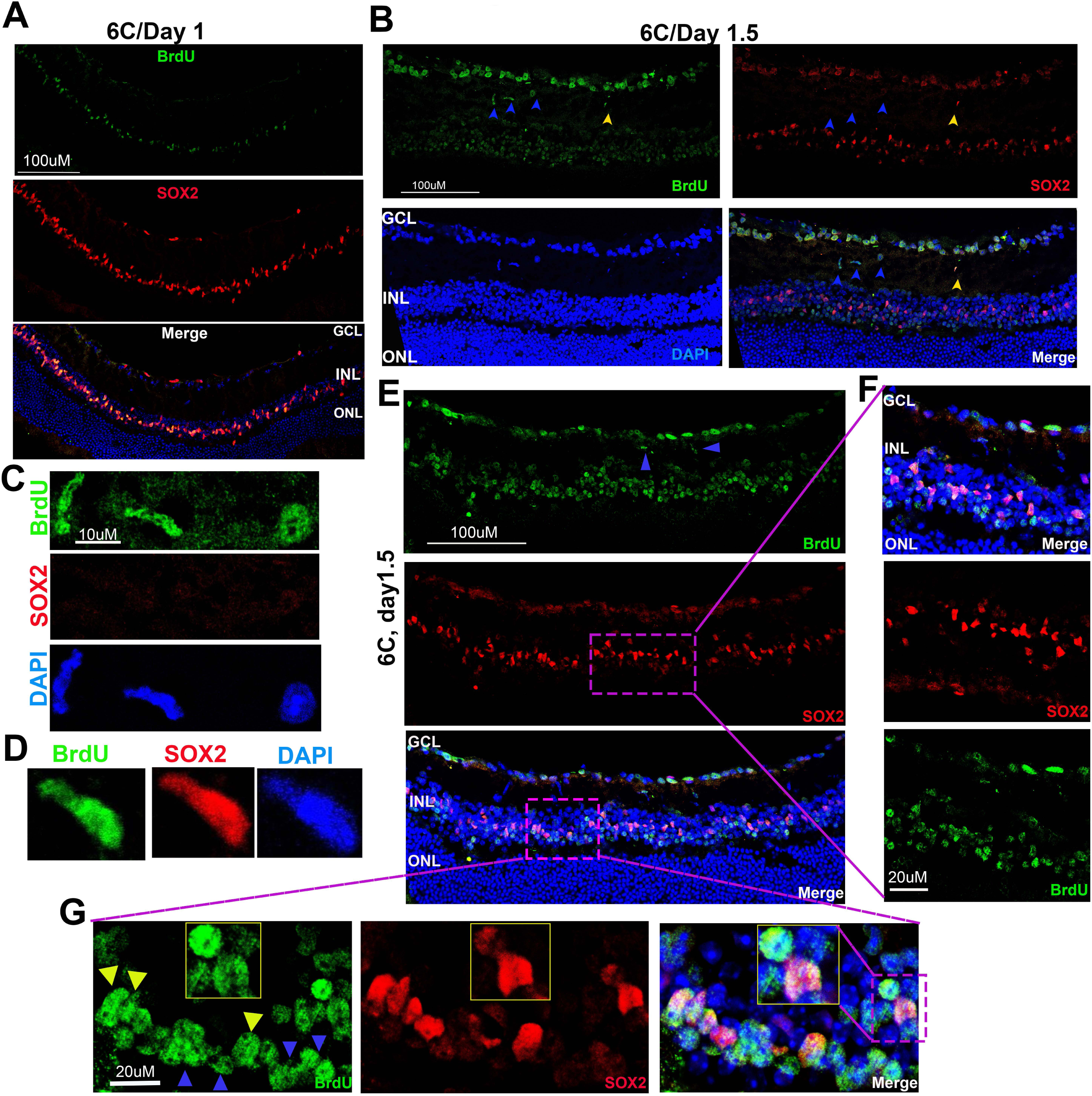
Generation and migration of BrdU^+^SOX2^-^ and BrdU^+^SOX2^+^ intermediate cells in the INL following 6C treatment in DBA/2J mice. **A.** On day 1, BrdU+ cells in the INL express the Müller glia marker SOX2. **B.** between days 1 and 2, BrdU+SOX2-(blue arrow) and BrdU+SOX2+ (yellow arrow) cells emerge in the INL and migrate to the GCL through the IPL. **C, D.** Magnified views of BrdU+SOX2- and BrdU+SOX2+ intermediate cells shown in B. **E.** High-magnification (40X) images illustrating BrdU+SOX2- and BrdU+SOX2+ cells migrating toward the GCL. **F, G.** Generation of BrdU+SOX2-cells in the INL (from dotted box in E, blue arrows).

**Figure S6.**
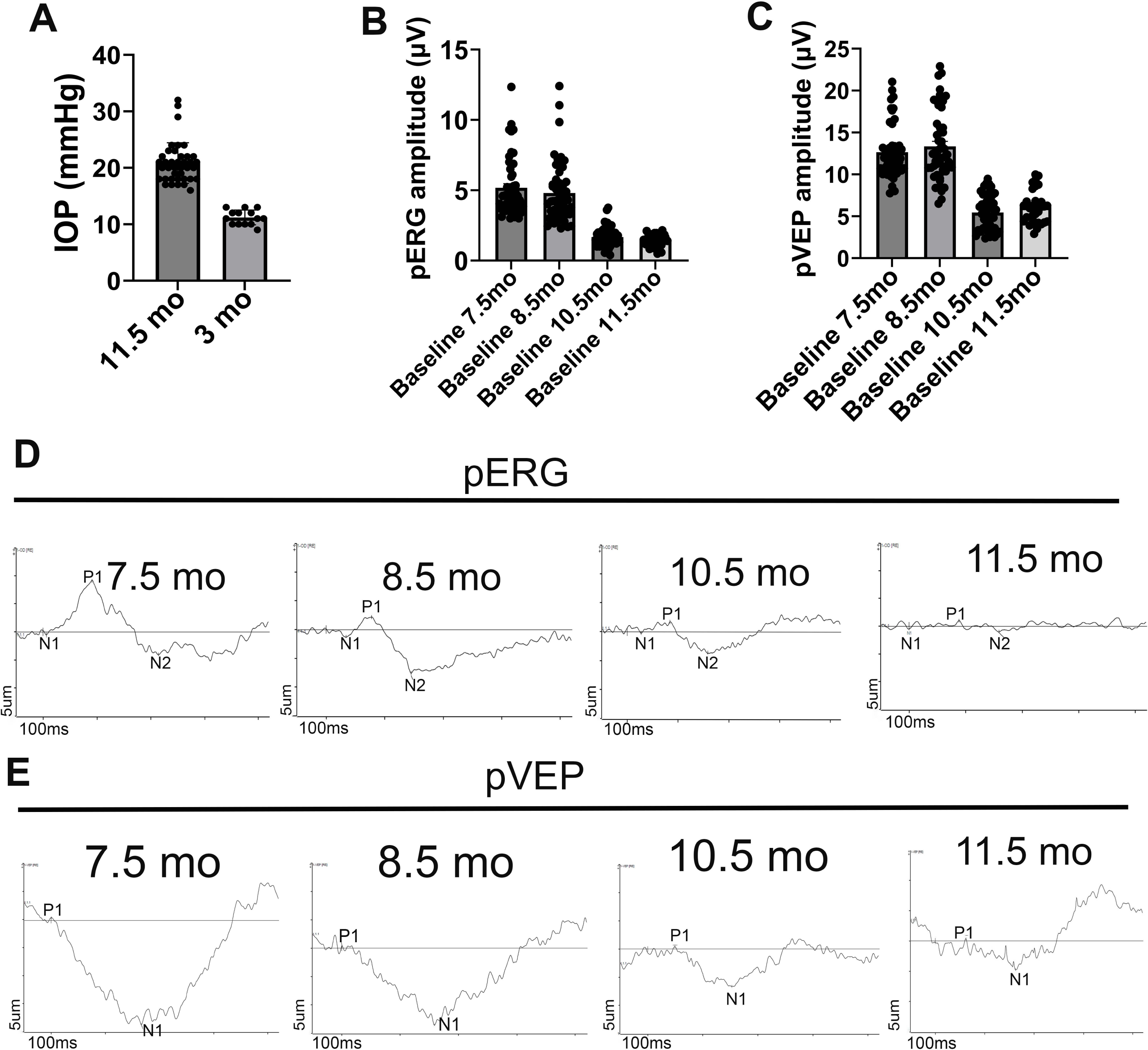
Generation of high intraocular pressure and longitudinal assessment of RGCs function in DBA/2J mice. **A.** Measurements of IOP at 3- and 11.5-months old mice. **B, C.** Decline in pERG and pVEP amplitudes from 7.5 months to 11.5 months DBA/2J mice**. D, E.** Representative pERG and pVEP waveforms at 7.5, 8.5, 10.5 and 11.5 months in DBA/2J mice.

**Figure S7.**
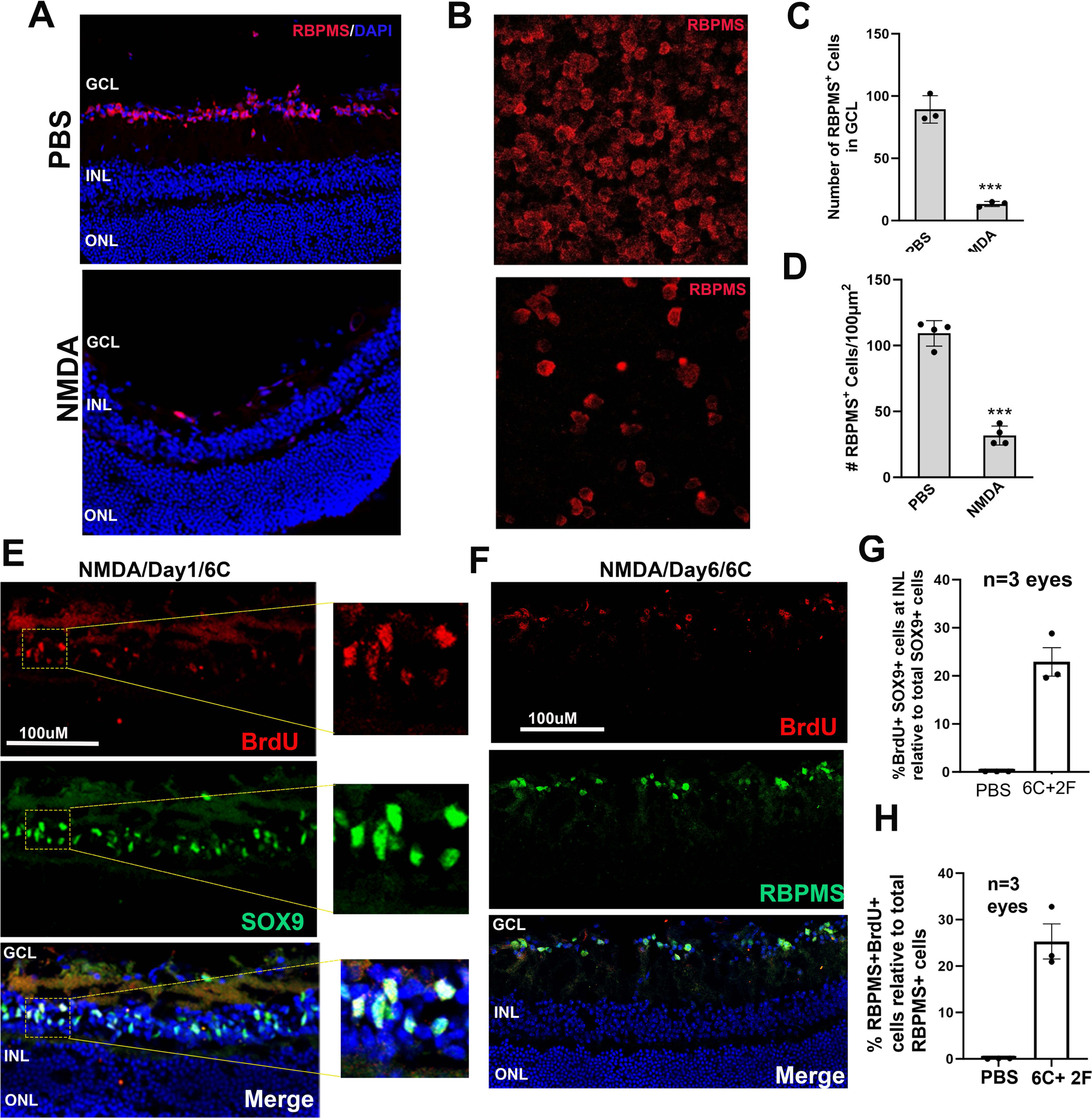
Preparation of NMDA injured RGC degeneration mouse model and *in vivo* CiGN reprogramming after 6C administration. A, **C.** Retinal sections stained with RBPMS in NMDA vs PBS treatment 7 days after injection. **B, D.** RBPMS+ cell count in GCL of PBS vs NMDA treated retina. **E, F.** Day1 and Day 3 retinal sections demonstrating BrdU+SOX9+ Muller glia cell cycle entry on day1 and subsequent reprogramming to CiGN cells by day6. **G, H.** BrdU^+^SOX2^+^ and BrdU^+^RBPMS^+^ cell count in the INL and GCL respectively after 6C treatment. n=3 eyes, 4 sections/eye, 3 fields/section were counted.

**Figure S8.**
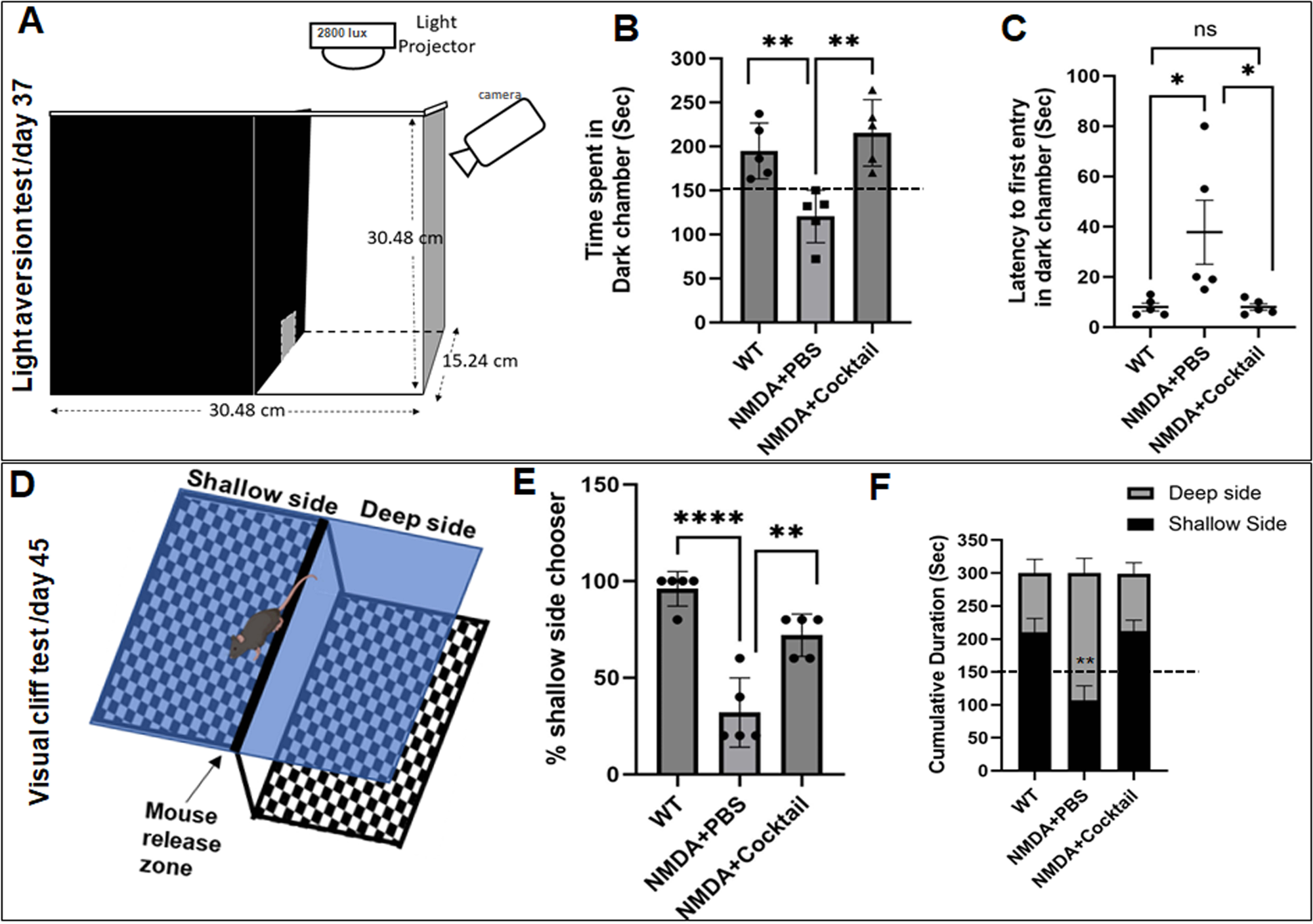
Rescue of light aversion & depth perception behavior after 6C treatment in NMDA injured C57/BL6 mice. **A.** Schematic of the experimental set up for light aversion behavior test. **B.** Plot of mean time (± SEM) spent in the dark chamber by the indicated experimental groups. **C.** Plot of mean latency time (± SEM) required to first entry to the dark chamber by the indicated experimental groups. **D.** Schematic of the experimental set up for death perception test (visual cliff test). **E.** Plot for percent shallow side chooser. Five trials were performed for each animal. **F.** Plot for cumulative duration spent deep or shallow sides by the indicated experimental groups after removal of the cliff. 1-Way ANOVA was used to determine significance in b, c, e. 2-Way ANOVA was used to determine significance in F.

**Figure S9.**
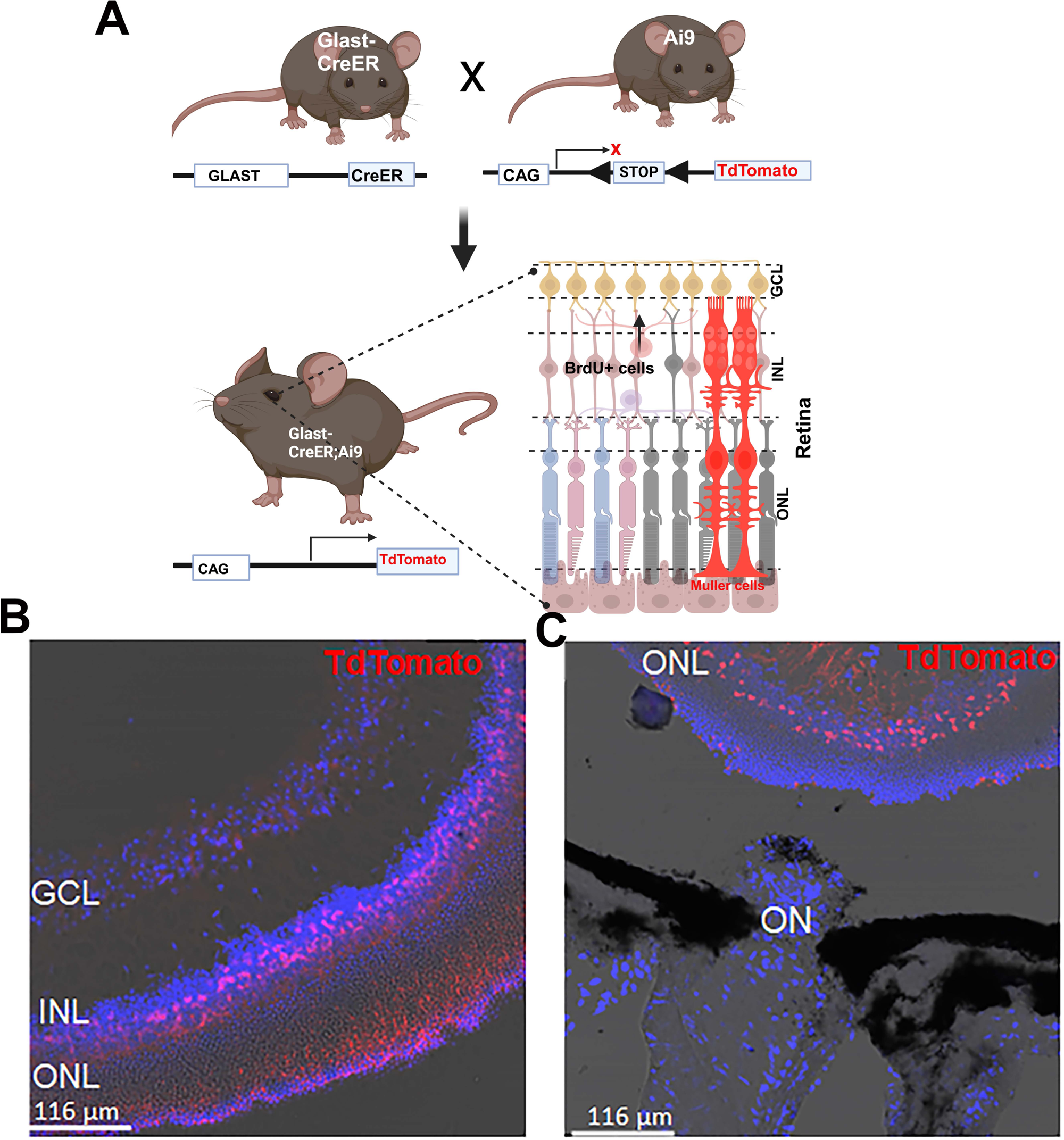
**A.** Schematics of Glast-CreER;Ai9 mice line generation. **B.** TdTomato expression in the Muller glia and no expression in GCL. **C.** No Tdtomato+ axons pass through the optic nerve head.

**Figure S10.**
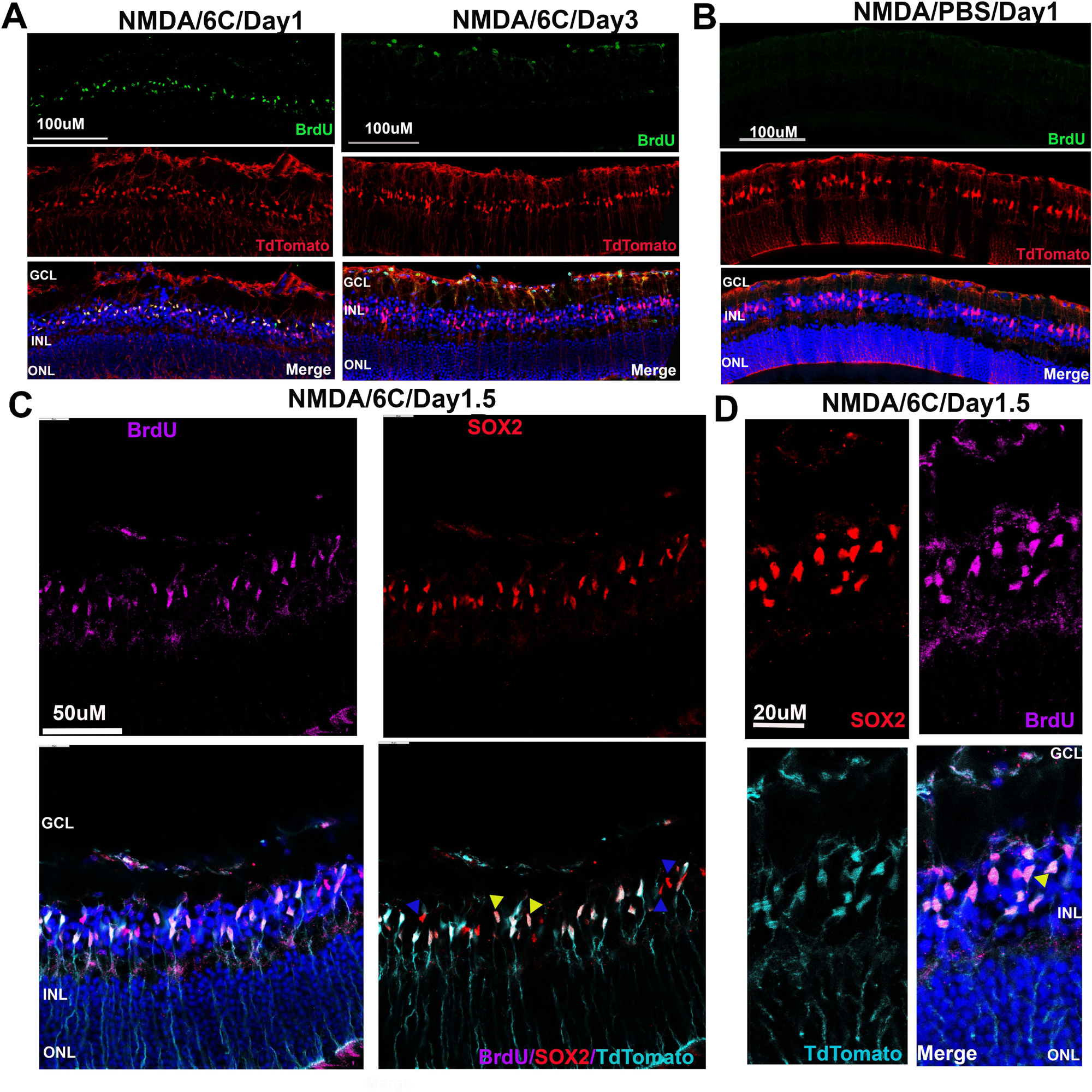
CiGN reprogramming in Glast-CreER;Ai9 reporter mice (post NMDA) occurs through an intermediate cellular stage possibly generated through asymmetric cytoplasmic divisions. A, B, Generation of BrdU^+^ cells in the INL on day1 and their subsequent migration to day3, similar to what we found in our DBA/2J and NMDA functional study. No BudU^+^ cells were evident in PBS injected control retina. **C**, Generation of SOX2^+^BrdU^+^TdTomato^+^ (yellow arrowhead) and SOX2^+^BrdU+TdTomato^-^ (blue arrowhead) intermediate cells in INL during 6C reprogramming. **D**, Images showing asymmetric divisions (yellow arrowhead) generate SOX2^+^BrdU^+^TdTomato^+^ and SOX2^+^BrdU^+^TdTomato^-^ cells in the INL on day 1.5 after 6C administration.

**Figure S11.**
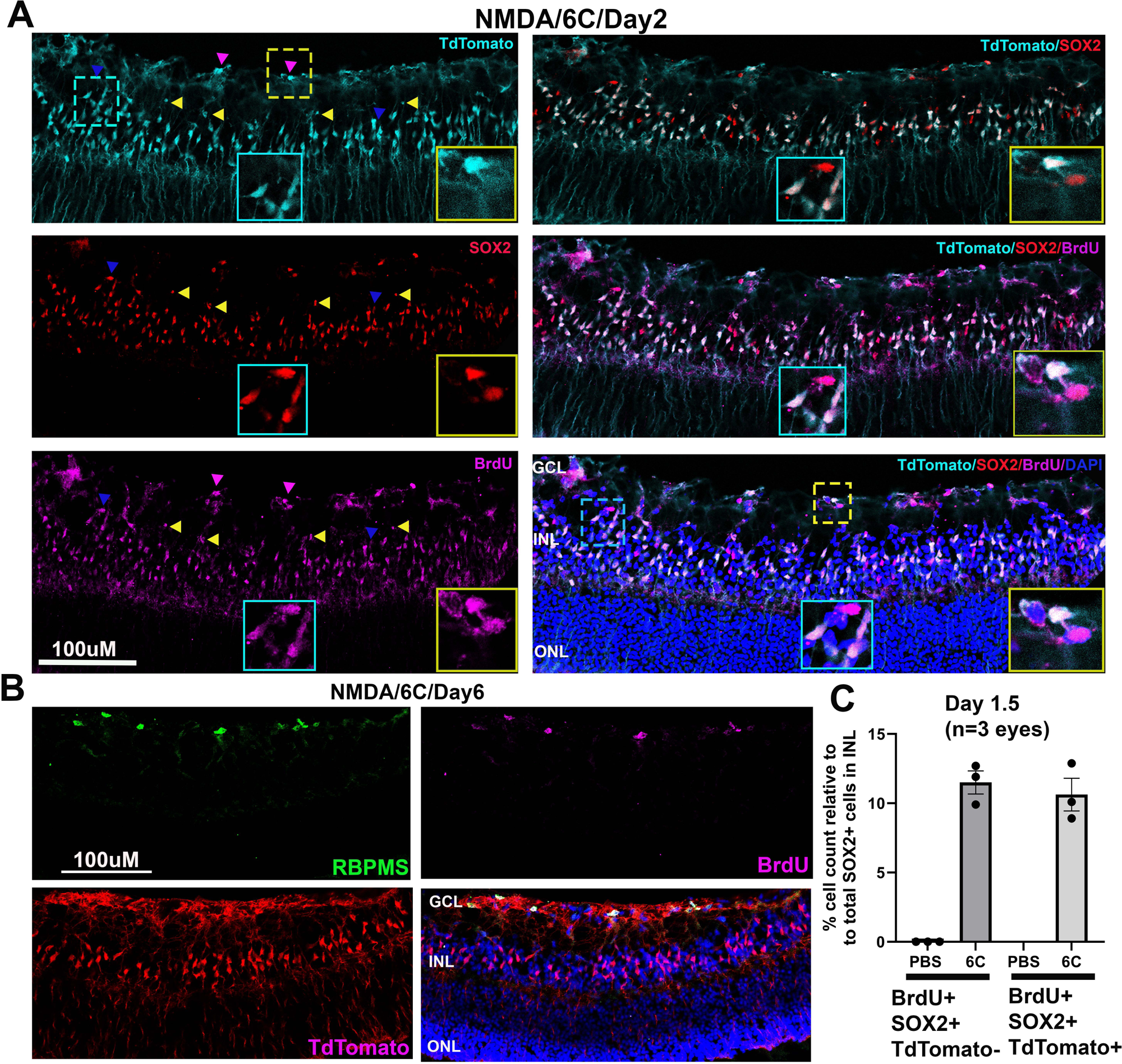
Generation and migration of intermediate cells to GCL after 6C reprogramming post NMDA in Glast-CreER;Ai9 mice. **A.** Migration of SOX2+BrdU+TdTomato+ (yellow arrowhead) and SOX2+BrdU+TdTomato-(blue arrowhead) cells from INL to GCL on day 2. Some cells have already reached GCL at this timepoint. Yellow boxes (corresponding to yellow dotted area) showing asymmetric divisions may also occur in GCL. Cyan boxes showing both migrating cell types on day 2. **B.** BrdU^+^ cells in GCL express RGC marker RGC on day6. **C.** cells count in the INL (n= 3 eyes, 4 sections/eye, 3 fields/section).

**Figure S12.**
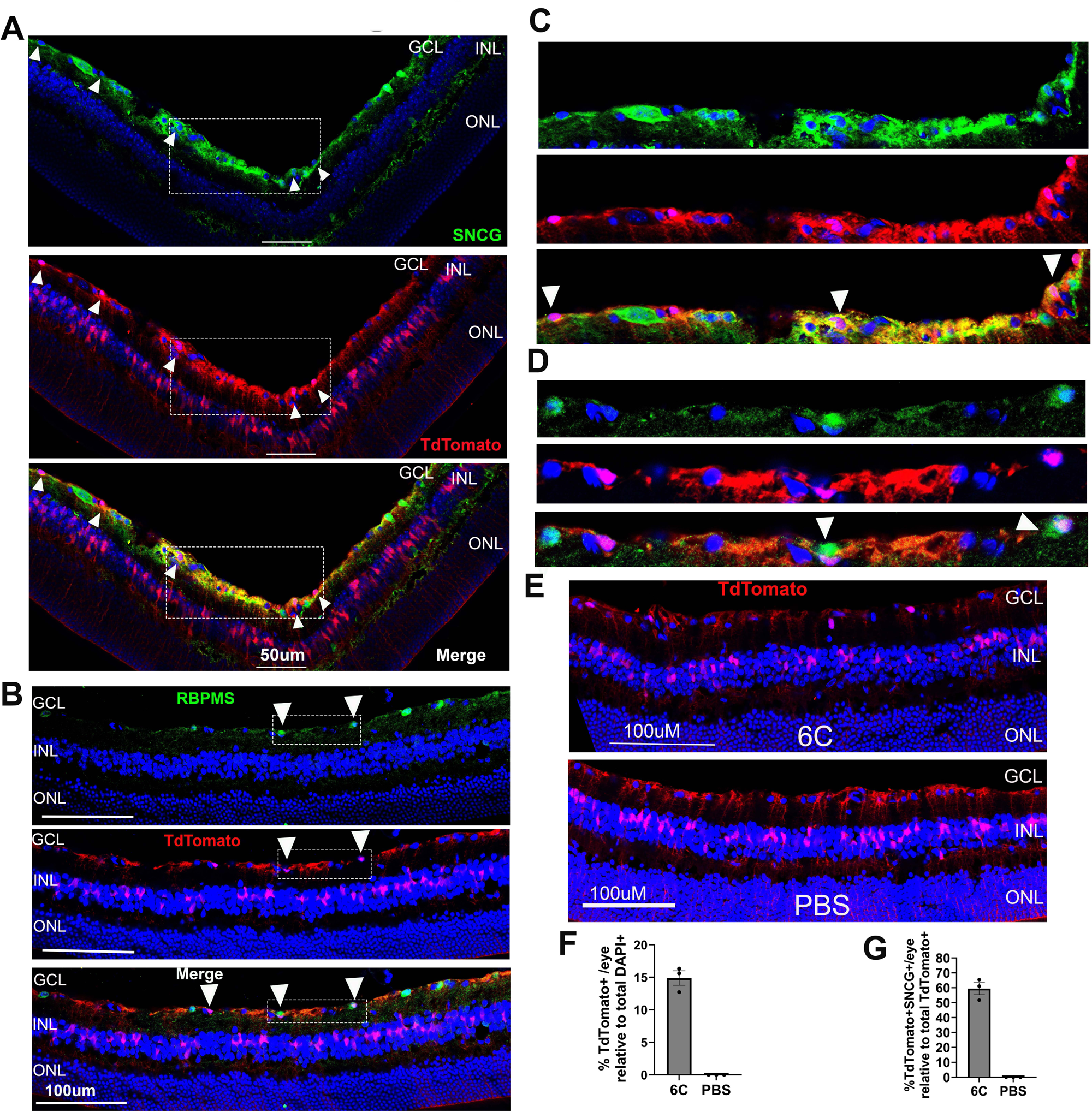
Lineage tracing in GLAST-Cre;Rosa26-TdTomato reporter mice after 6C treatment in NMDA injured mice eye. A, **B.** TdTomato^+^ cells in GCL are positive for SNCG and RBPMS indicating their Müller cell origin and reprogramming to RGCs. **C, D.** Magnified images showing TdTomato^+^/SNCG^+^ and TdTomato^+^/RBPMS^+^ cells (white arrows). TdTomato expression was in the nucleus. **E.** PBS control did not show the nuclear expression of TdTomato in the GCL post 6C. **F, G.** Percent TdTomato^+^ and TdTomato^+^/SNCG^+^ cell count in 6C treated eye. n=3 eyes, 4 sections/eye, 3 fields/section were counted. n=3 eyes examined.

**Figure S13.**
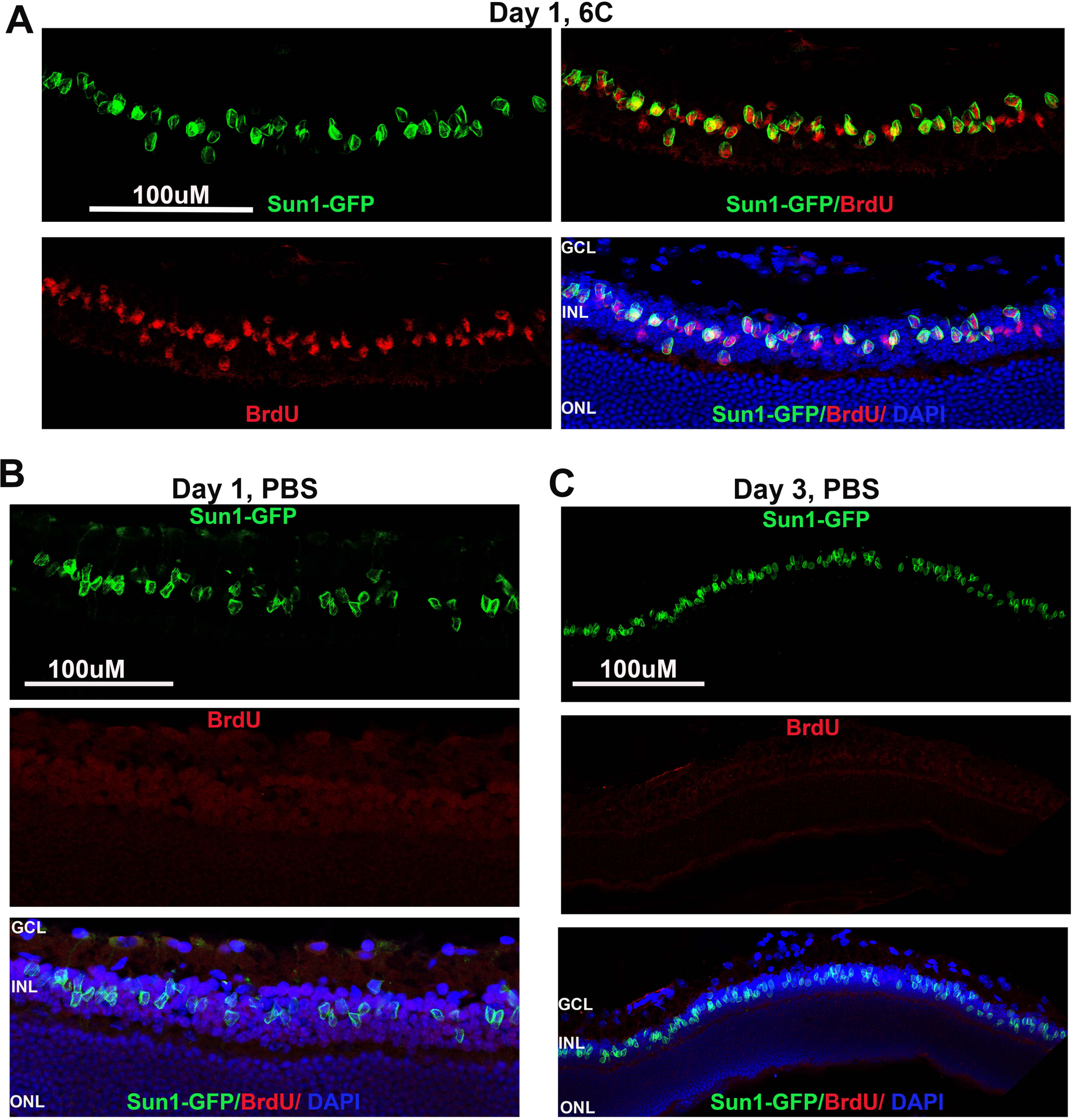
Muller glia cell cycle entry in Glast-Cre:Sun1-GFP mice post NMDA. **A.** 6C treated retina showing GFP^+^BrdU^+^ cells in the INL on day1. **B, C.** No BrdU+ cells were evident in PBS injected retina on days 1 and 3.

**Figure S14.**
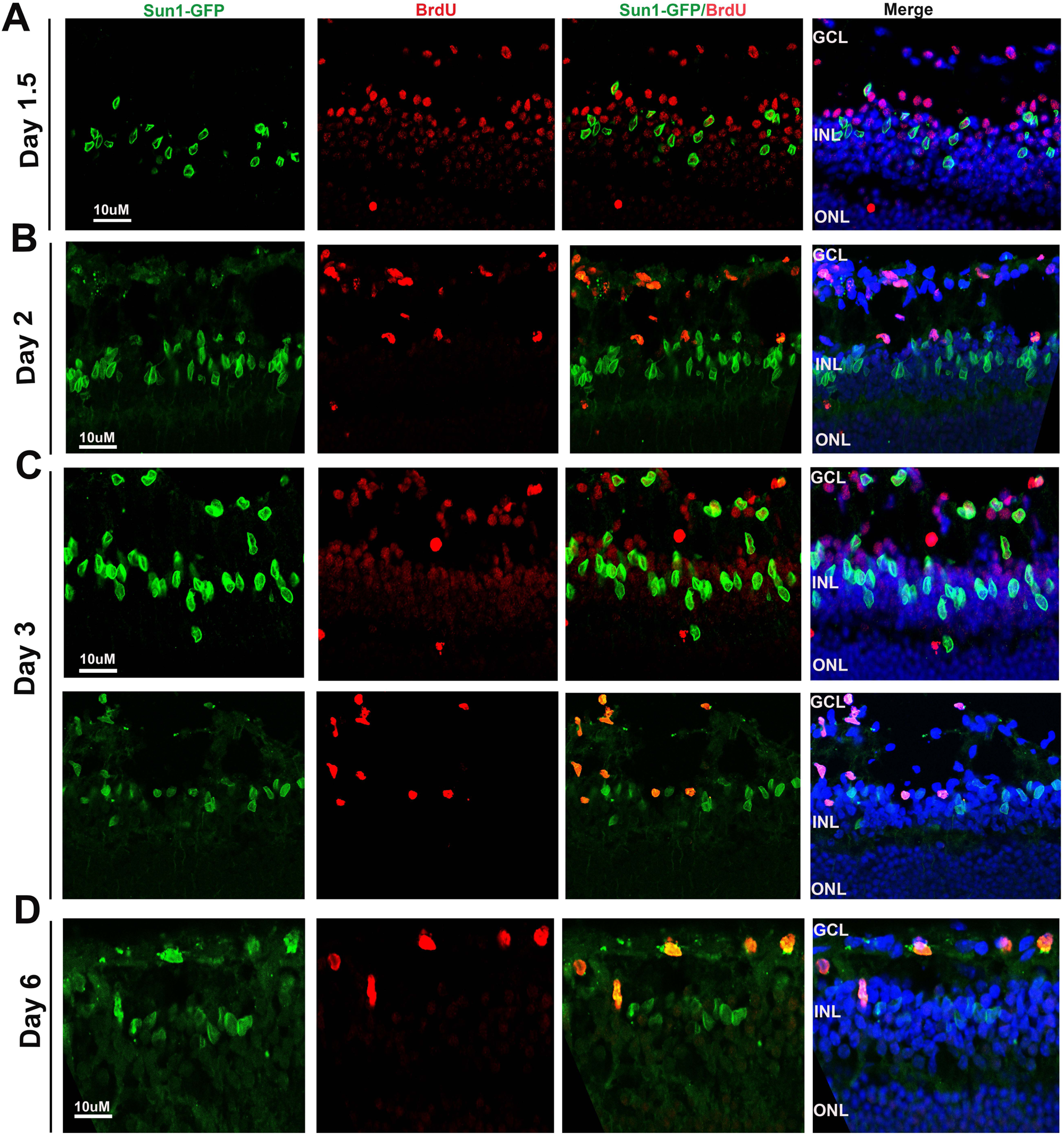
Migration of GFP^+^BrdU^+^ and GFP^-^BrdU^+^ cells from INL to GCL upon 6C treatment post NMDA in Glast-Cre:Sun1-GFP mice. **A.** GFP^+^BrdU^+^ and GFP^-^BrdU^+^ cells migratory cells in the IPL on day 1.5. **B.** On day 2 many GFP^+^BrdU^+^ and GFP^-^BrdU^+^ cells reached GCL. **C.** GFP^+^BrdU^+^ cells are divided (based on morphology) after reaching GCL. **D.** On day 3, GFP^+^BrdU^+^ in the GCL remains in the GCL for possible differentiation.

**Figure S15.**
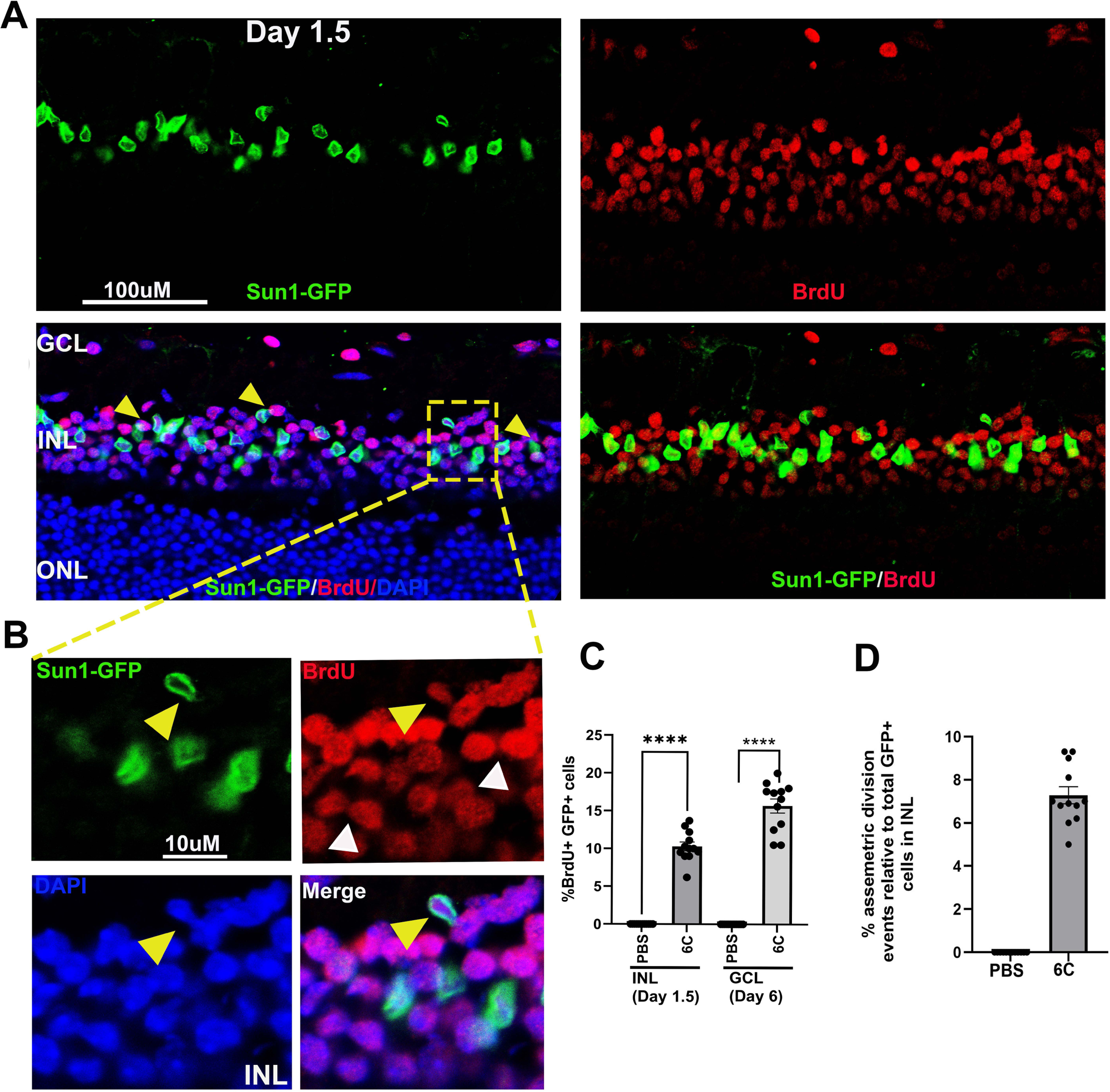
Generation of BrdU^+^ intermediate cells in the INL likely occurs through asymmetric divisions in Glast-Cre:Sun1-GFP mice post NMDA. **A.** Retina from 6C treated GFP^+^BrdU^+^ cells in the INL on day 1.5. yellow arrowheads in the merged image showing asymmetric divisions. **B.** Enlarged image showing asymmetric divisions occurs in the INL during CiGN reprogramming (yellow arrowhead) followed by BrdU+ cell proliferations (white arrowheads). **C, D.** quantification of GFP+BrdU+ cells in the INL (day 1.5) and GCL (day 6) and number of asymmetric division events in the INL of day 1.5. n=3 eyes. 4 sections/eye, 3 fields/section was counted.

**Figure S16.**
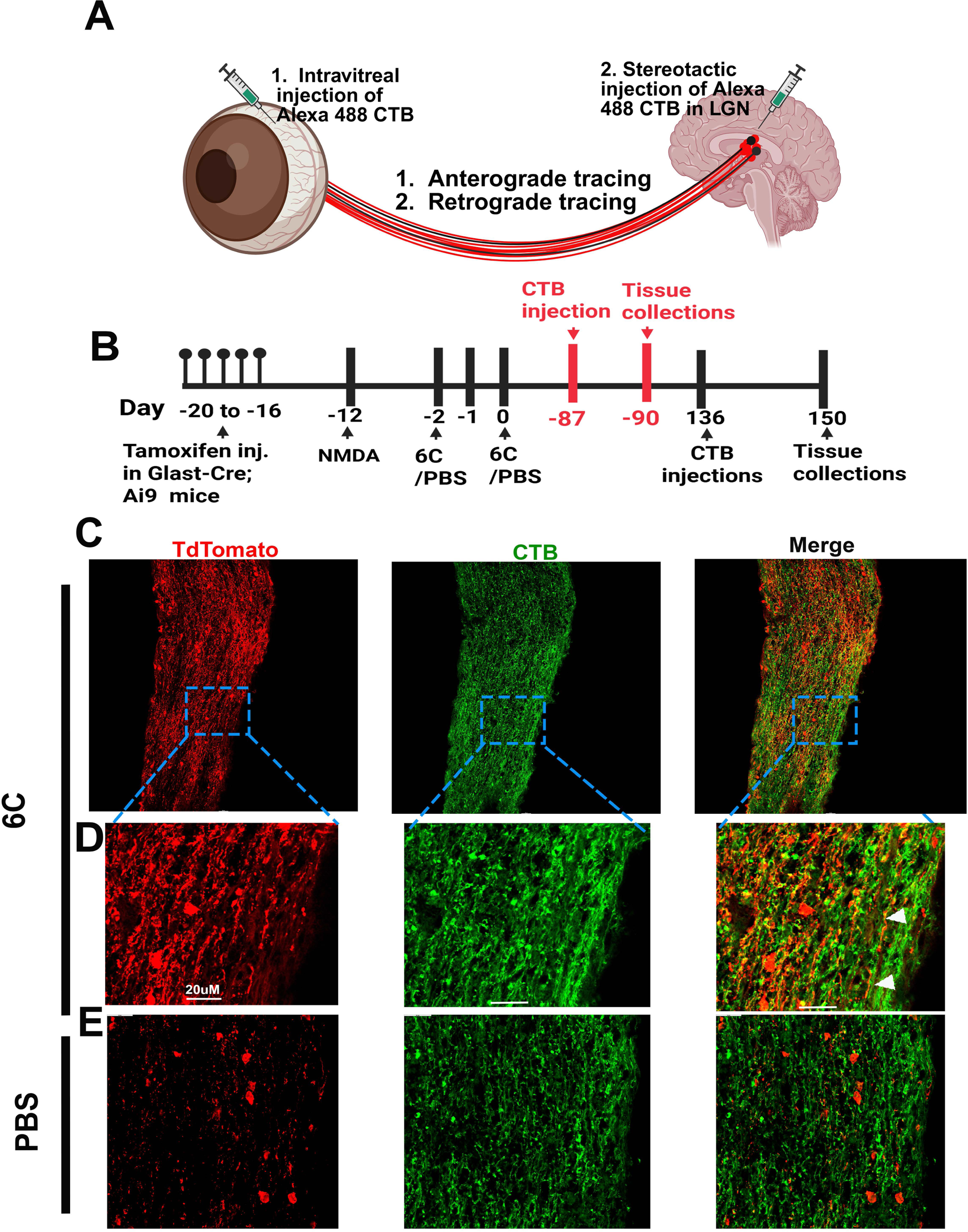
Transport of Cholera Toxin B (CTB-488) through TdTomato+ CiGN axons on day 90. **A.** Schematic diagram of anterograde and retrograde tracing experiment. **B.** Experimental timeline for anterograde and retrograde tracing experiments in Glast0Cre;Ai9 mice. **C.** TdTomato+ axons co-labeled with CTB staining (25X image). **D.** Magnified view of the indicated regions. CTB was injected three days before the experimental endpoint. **E.** Optic nerves from PBS injected control. CTB+/TdTomato+ axons are indicated by white arrowheads. n = 3 optic nerves were analyzed.

**Figure S17.**
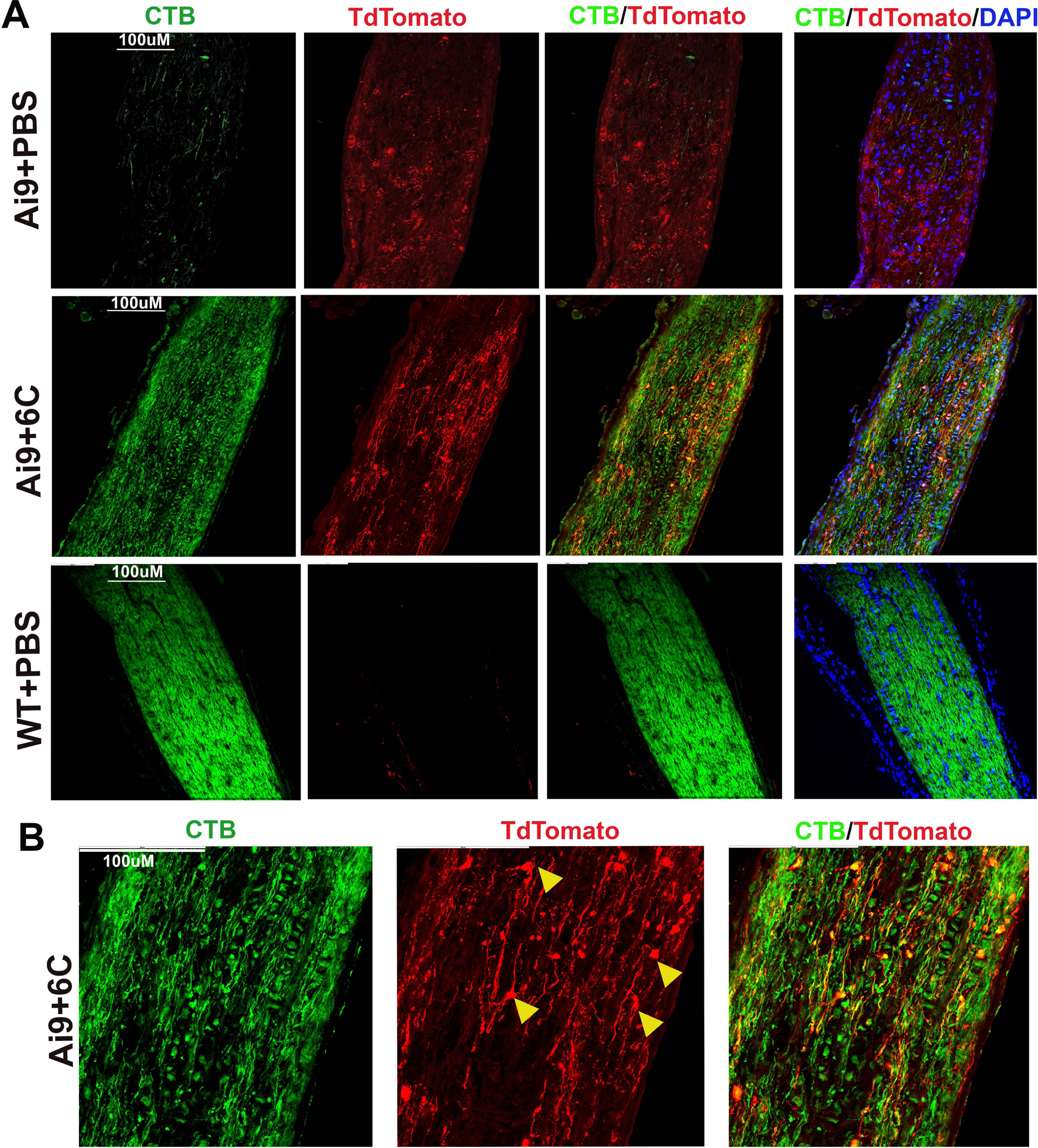
Retrograde transport of Cholera Toxin B (CTB-488) through TdTomato+ CiGN optic nerve axons on day 150. **A.** CTB transport in PBS and 6C treated Ai9 mice. PBS injected wildtype C57/BL6 optic nerve as control. **B.** 40X magnification of optic nerve from 6C treated Glast-Cre:Ai9 mice. Prospective bridge neurons are indicated in yellow arrowheads.

**Figure S18.**
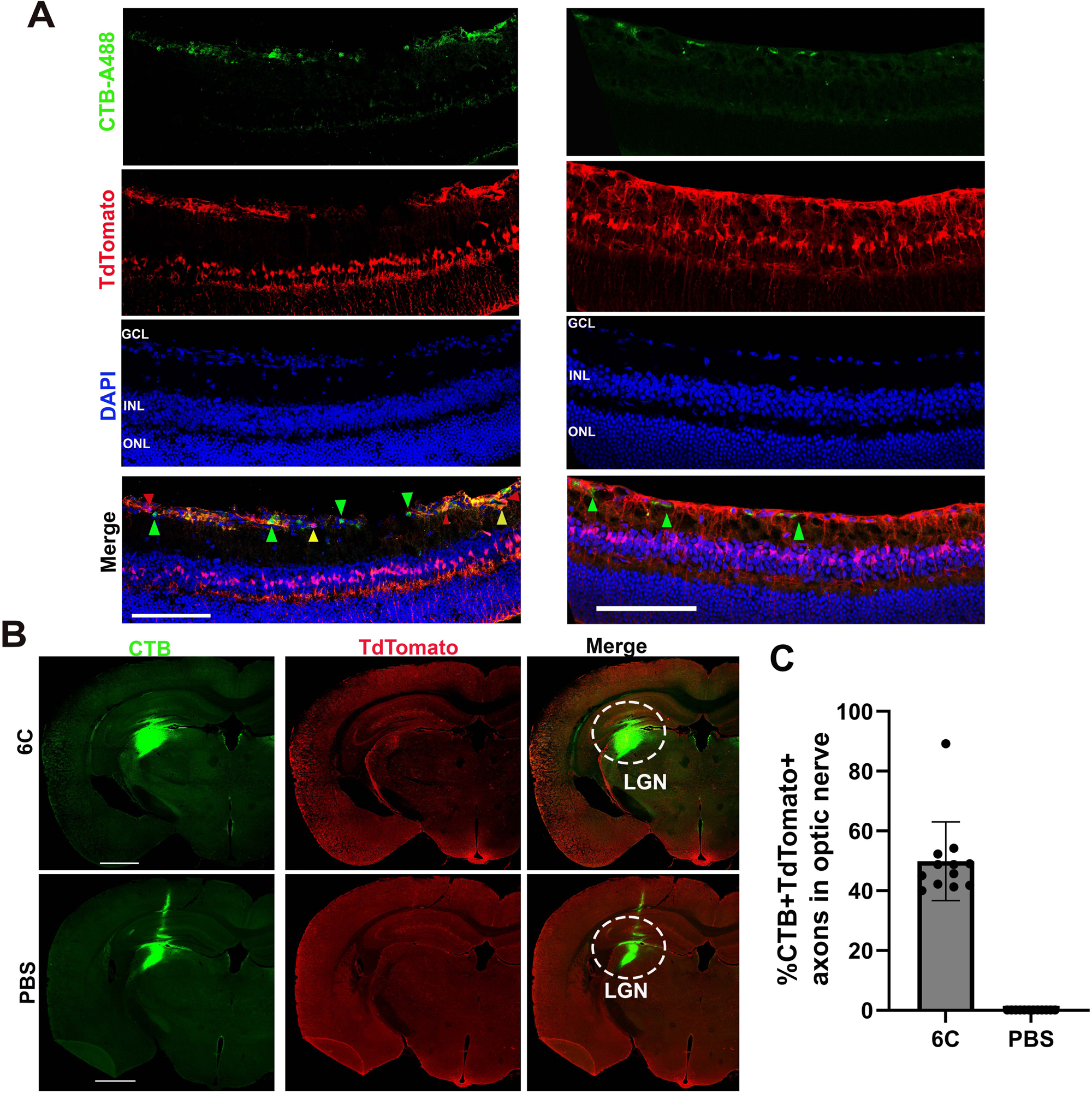
Retrograde transport of Cholera Toxin B (CTB-488) in Glast-Cre:Ai9 retina on day 150. **A.** Three different types of cells were evident: (i) CTB+TdTomato-(host RGCs), CTB+TdTomato+ (possible regenerated CiGN), CTB-TdTomato+ regenerated CiGN those did not reach to the LGN. **B.** CTB injection sites in both treatments. **B.** Quantification for the number of CTB+TdTomato+ axons in the optic nerve (n=3 optic nerve counted, 4 sections/optic nerve).

**Figure S19.**
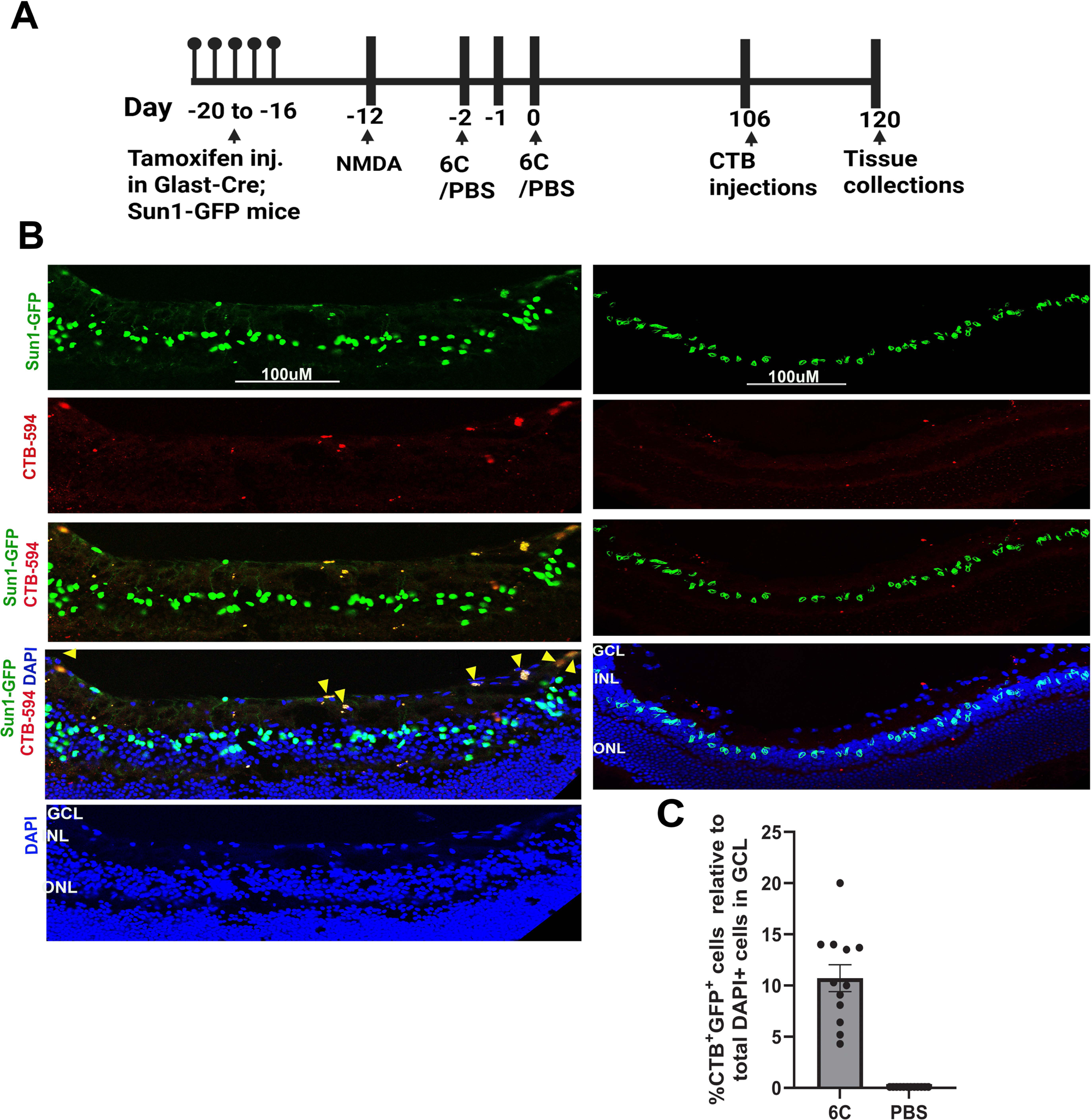
Retrograde transport of Cholera Toxin B (CTB-594) in Glast-Cre:Sun1-GFP retina on day 150. **A.** Experimental time line for anterograde and retrograde tracing experiments in Glast-Cre;Sun1-GFP mice. **B.** Presence of GFP+ and CTB+ cells in the GCL indicating CiGN (GFP+ here) established connections to the LGN. **C.** Quantification for the number of CTB+GFP+ cells in GCL (n=3 eyes, 4 sections/eye, 3 fields/section counted).

**Figure S20.**
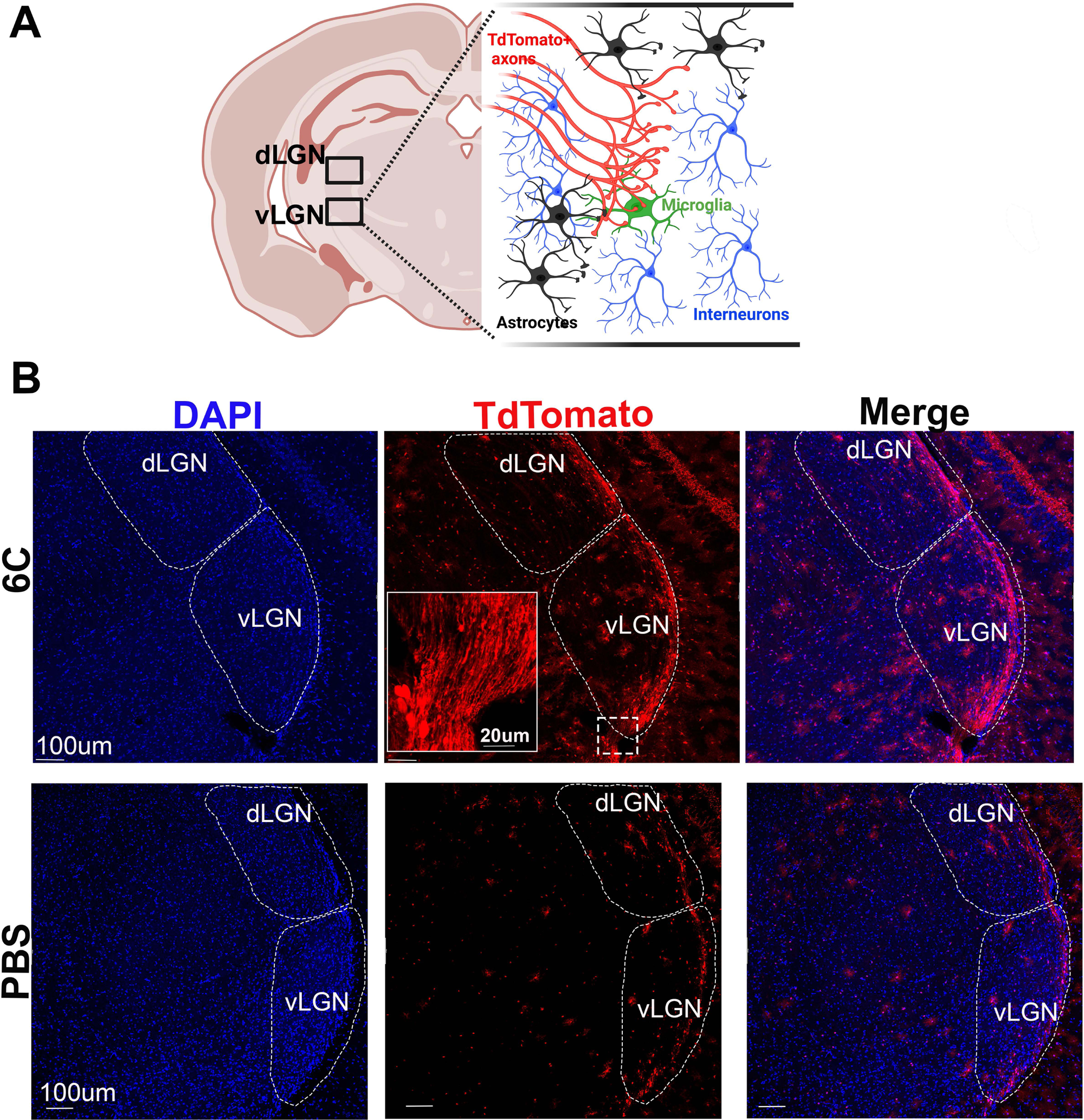
*In vivo* reprogrammed CiGN cells establish connections to the LGN in 6C treated GLAST-Cre;Ai9 reporter mice. **A.** Schematic of TdTomato+ axon projections in dorsal lateral geniculate nucleus (dLGN) and ventral geniculate nucleus (vLGN). **B.** Images of LGN showed dLGN and vLGN contains TdTomato+ axons projections in 6C treated mice. Inset upper middle panel showing TdTomato+ neuronal projections entering vLGN in 6C treatment. TdTomato+ axons are absent in enlarged dLGN and vLGN of PBS treated mice.

**Figure S21.**
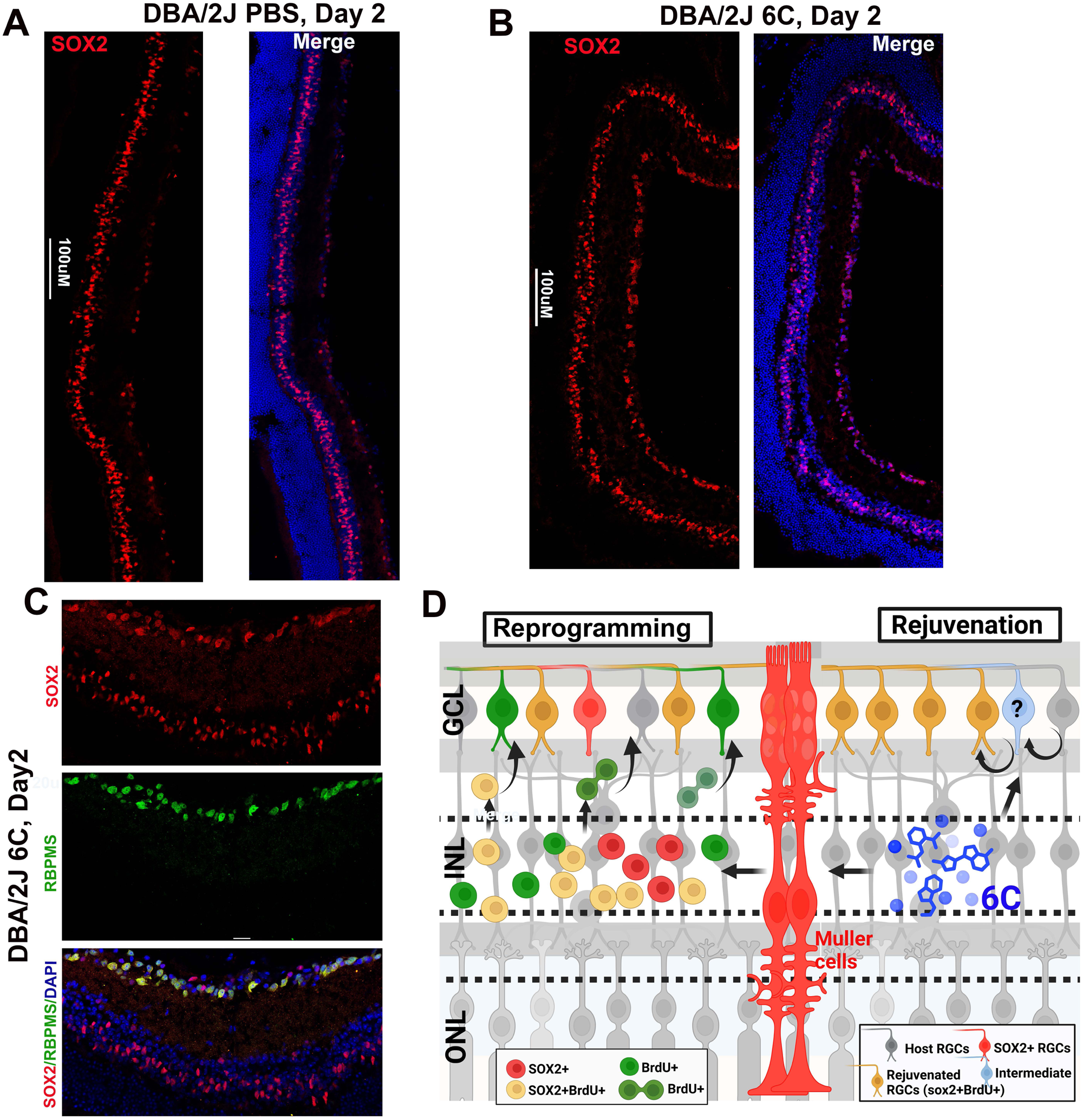
Enhanced SOX2 expression following 6C treatment may contribute to vision restoration in DBA/2J mice. A, **B.** SOX2 expression in the retina of PBS- and 6C-treated mice between days 1 and 2 after 6C injection. **C.** SOX2-expressing cells in the GCL co-express RBPMS, indicating their identity as RGCs. **D.** Mechanism of CiGN generation by 6C treatment and their proposed role in vision restoration (schematic presentation).

**Figure S22.**
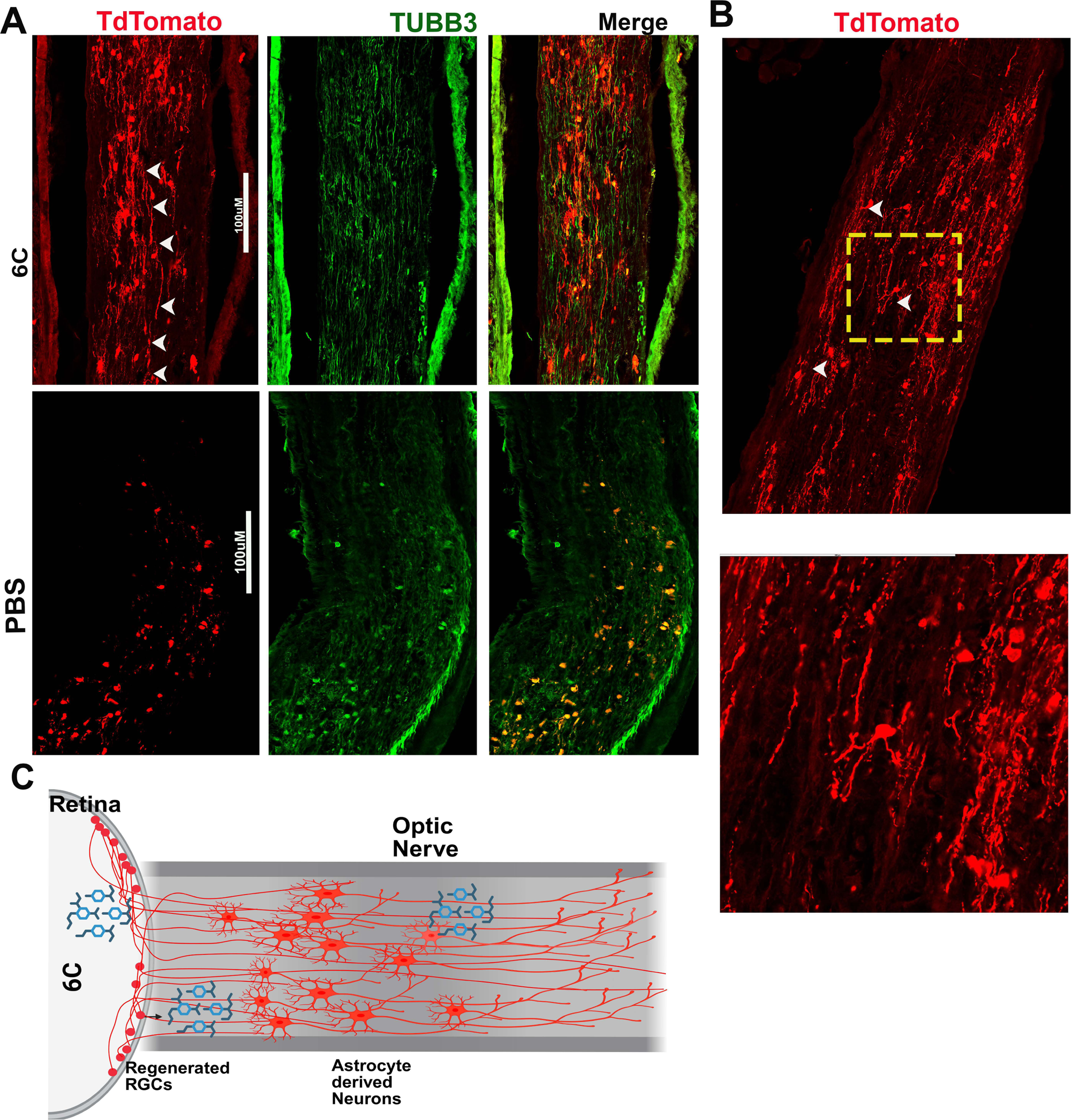
Formation of neuronal relays between regenerated CiGN axons and optic nerve astrocyte–derived neurons may serve as a potential mechanism for re-establishing eye-to-brain connectivity. **A.** 6C and PBS treated optic nerve from Glast-Cre;Ai9 mice 120 days after 6C injections. **B.** Enlarged images showing an event of neuronal relay formation. **C.** Schematic presentation of neuronal relay formation.

## Notes

### Competing Interest Statement

The authors have declared no competing interest.

### Summary of Updates

Loss of retinal ganglion cells (RGCs) is a major cause of vision loss in optic neuropathies such as glaucoma, with no available treatments to restore vision. In teleost fish, Muller glia possesses a remarkable regenerative capacity to replace lost RGCs and restore vision a capability lacking in mammals. Here, we have identified a six-small molecule cocktail (6C) that induces in vivo reprogramming of retina resident Muller glia into retinal neurons within the ganglion cell layer (GCL) following RGC injury. We name these cells chemically induced GCL neurons (CiGN). During reprogramming process, Muller glia re-enters the cell cycle in the inner nuclear layer, asymmetrically divide, proliferate and migrate to the GCL as SOX2+ and SOX2− intermediates, exit the cell cycle, and differentiate into CiGN cells mirroring some aspects of retinal regeneration seen in teleost fish. Functionally, 6C treatment restores long-term visual functions in rodent models of ocular hypertension and NMDA-induced RGC injury. Notably, 6C induces axon extension along the optic nerve and establish connections to the lateral geniculate nucleus (LGN) possibly through a neuronal relay mechanism. These findings highlight small molecule mediated cellular reprogramming as a potential therapeutic strategy for vision restoration in glaucoma and other optic neuropathies that affects millions of children and adults worldwide.

