## Supplemental Tables 1-6 for "Chemically induced *in vivo* retinal neuron regeneration rescues vision in mice" (from another version of this preprint: version 1, 2023-12-27)

**Supplementary table 1:** Small molecules stock and final concentration for *in vivo* CiRGC reprogramming.

<table>
<tr><td>Molecules</td><td>Stock concentration</td><td>Concentration of test article in injection cocktail (50 µL)</td><td rowspan="11">1.5 µl from the cocktail was injected in each eye</td></tr>
<tr><td>VPA (V)</td><td>250 mM</td><td>10 mM</td></tr>
<tr><td>CHIR (C)</td><td>12 mM</td><td>96 µM</td></tr>
<tr><td>RepSox (R)</td><td>20 mM</td><td>40 µM</td></tr>
<tr><td>Forskolin (F)</td><td>50 mM</td><td>200 µM</td></tr>
<tr><td>IWR1 (I)</td><td>10 mM</td><td>200 µM</td></tr>
<tr><td>Noggin</td><td>20 µg/mL</td><td>800 ng/mL</td></tr>
<tr><td>IGF-1</td><td>200 µg/mL</td><td>8 µg/mL</td></tr>
<tr><td>S24795</td><td>50 mM</td><td>1 mM</td></tr>
<tr><td>BrdU IVT</td><td>10 mg/mL</td><td>1 mg/mL</td></tr>
<tr><td>BrdU IP</td><td>10 mg/mL</td><td>50mg/kg</td></tr>
</table>

**Supplementary table 2:** Small molecule stock concentrations and volume for preparing a 50 µL solution cocktail.

| *Molecules* | *Stock concentration* | *Amount added in 50 µL total volume* |
|---|---|---|
| VPA (V) | 250 mM | 2 µL |
| CHIR (C) | 12 mM | 0.4 µL |
| RepSox (R) | 20 mM | 0.1 µL |
| Forskolin (F) | 50 mM | 0.2 µL |
| IWR1 (I) | 10 mM | 1 µL |
| Noggin | 20 µg/mL | 2 µL |
| IGF-1 | 200 µg/mL | 2 µL |
| S24795 | 50 mM | 1 µL |
| BrdU | 10 mg/mL | 5 µL |
| PBS | 10 mg/mL | To 50 µL |

**Supplementary table 3:** Retinal Differentiation Medium 2 (RDM2) Composition

| **CiRGC Differentiation Medium** | | | |
|---|---|---|---|
| **Components** | **Stock Conc.** | **Final Conc.** | **Volume added in 50 mL** |
| Knockout Serum Replacement | 100 % (regular stock) | 10% | 5 ml |
| B27 Supplement | 100x | 1% | 500 µl |
| N2 Supplement | 100x | 1% | 500 µl |
| Noggin | 20 µg/ml | 10 ng/ml | 25 µl |
| DKK1 | 10 µg/ml | 10 ng/ml | 50 µl |
| IGF-1 | 200 µg/ml | 50 ng/ml | 12.5 µl |
| bFGF | 100 µg/ml | 5 ng/ml | 2.5 µl |
| SB431542 | 50 mM | 5 µM | 5 µl |
| DMEM/F12 with Glutamax | | | Up to 50 ml |
| **RDM1 medium composition** | | | |
| RDM2 minus SB431542 | | | |

**Supplementary table 4:** Small molecules stock and final concentration for *in vitro* CiRGCs reprogramming.

| **Small molecules/Growth Factor** | **Stock concentration** | **Final amount used, ul/ml of medium** | **Final conc.** | **Source, cat #** |
|---|---|---|---|---|
| VPA (V) | 250mM | 2 | 0.5mM | Sigma, P4543 |
| CHIR (C) | 12mM | 0.4 | 4.8 µM | Cayman, 13122 |
| Repsox (R) | 20mM | 0.1 | 2 µM | Cayman, 14794 |
| Forskolin (F) | 50mM | 0.2 | 10 µM | Cayman,11018 |
| ISX9 (Is) | 50 mM | 0.4 | 20 µM | Tocris, 4439 |
| Sonic Hedgehog (Shh) | 2.5 µM | 1.2 | 3nM | R&D System,1314sh |
| Taurine (T) | 100mM | 1 | 100 µM | Sigma, T0625 |

| | | | | |
|---|---|---|---|---|
| Retinoic Acid (R) | 1 mM | 1 | 1 µM | Sigma, R2615 |
| Noggin | 20 µg/ml | - | 10 ng/ml | PeproTech,120-10C |
| IGFI | 200 µg/ml | - | 50 ng/ml | PeproTech,100-11 |

**Supplementary table 5:** Primers for RT-qPCR

| Species | Gene | Forward | Reverse |
|---|---|---|---|
| Human | Brn3a | GGGCAAGAGCCATCCTTTCAA | CTGTTCATCGTGTGGTACGTG |
| Human | Brn3b | CAAGCAGCGACGCATCAAG | GGGTTTGAGCGCGATCATATT |
| Human | Sncg | TGAGCAGCGTCAACACTGTG | GAGGTGACCGCGATGTTCTC |
| Human | Isl-1 | TACGGGATCAAATGCGCCAA | CACACAGCGGAAACACTCGAT |
| Human | Nef-H | GCAGTCCGAGGAGTGGTTC | CGCATAGCGTCTGTGTTCA |
| Human | Nef-L | ATGAGTTCCTTCAGCTACGAGC | CTGGGCATCAACGATCCAGA |
| Human | Beta III Tubulin | GGCCAAGGGTCACTACACG | GCAGTCGCAGTTTTCACACTC |
| Hyman | GAPDH | AAGGTGAAGGTCGGAGTCAAC | GGGGTCATTGATGGCAACAATA |

**Supplementary table 6:** Antibodies used for this study.

| Name of Antibody | Host | Dilution | Vendor | Cat# |
|---|---|---|---|---|
| Anti-Brn3a | Mouse | 1:100 | Santa Cruz | SC-8429 |
| Anti-RBPMS | Rabbit | 1:500 | GeneTex | GTX118619 |
| Anti-B-tubulin III | Mouse | 1:1000 | Biolegened | MMS-435P |
| Anti-SNCG | Mouse | 1:500 | Abnova | H00006623-M01 |
| Anti-GFP | chicken | 1:1000 | Abcam | Ab13970 |
| Anti-Glutamine synthase | Mouse | 1:1000 | Invitrogen | MA5-27749 |
| Anti-Brdu | Rabbit | 1:500 | GeneTex | GTX128091 |
| Anti-TdTomato | Goat | 1:500 | SICGEN | AB818-200 |
